# Gaze onsets during naturalistic infant-caregiver interaction associate with ‘sender’ but not ‘receiver’ neural responses, and do not lead to changes in inter-brain synchrony

**DOI:** 10.1101/2022.05.27.493545

**Authors:** I. Marriott Haresign, E.A.M Phillips, M. Whitehorn, F. Lamagna, M. Eliano, L. Goupil, E.J.H. Jones, S.V. Wass

**Author notes:** Corresponding author – Mr I Marriott Haresign, University of East London, London E15 4LZ.

## Abstract

Temporal coordination during infant-caregiver social interaction is thought to be crucial for supporting early language acquisition and cognitive development. Despite a growing prevalence of theories suggesting that increased inter-brain synchrony associates with many key aspects of social interactions such as mutual gaze, little is known about how this arises during development. Here, we investigated the role of mutual gaze *onsets* as a potential driver of inter-brain synchrony. We extracted dual EEG activity around naturally occurring gaze onsets during infant-caregiver social interactions in N=55 dyads (mean age 12 months). We differentiated between two types of gaze onset, depending on each partners’ role. ‘Sender’ gaze onsets were defined at a time when either the adult or the infant made a gaze shift towards their partner at a time when their partner was either already looking at them (mutual) or not looking at them (non-mutual). ‘Receiver’ gaze onsets were defined at a time when their partner made a gaze shift towards them at a time when either the adult or the infant was already looking at their partner (mutual) or not (non-mutual). Contrary to our hypothesis we found that, during a naturalistic interaction, both mutual and non-mutual gaze onsets were associated with changes in the sender, but not the receiver’s brain activity and were not associated with increases in inter-brain synchrony above baseline. Further, we found that mutual, compared to non-mutual gaze onsets were not associated with increased inter brain synchrony. Overall, our results suggest that the effects of mutual gaze are strongest at the intra-brain level, in the ‘sender’ but not the ‘receiver’ of the mutual gaze.

## 1. Introduction

Most of our early life is spent in the presence of an adult social partner. Most early attention – and, in particular, most early cognitive learning – takes place in social settings (Csibra and Gergely, 2009). But almost all of our understanding of how the brain subserves attention and learning has come from studies that measure individual brains in isolation.

In recent years our understanding of early social and communicative development has relied heavily on studying ostensive signals, defined as signals from a communicator to generate an interpretation of communicative intention in an addressee. It has been argued that, from shortly after birth, infants’ brains are sensitive to ostensive signals (such as direct gaze, smiles and infant-directed vocalisations), and that ‘sender’ communicative signals play a key role in supporting early learning exchanges (Werker and Yeung, 2005; Csibra and Gergely, 2009; Csibra, 2010; Southgate et al., 2010; Begus et al., 2014; 2016; Ferguson & Lew-Williams, 2016). In this paper we focus on mutual gaze, which is a widely studied ostensive signal.

Farroni et al., 2002 found that images of faces showing direct vs averted eye contact elicited greater amplitude event related potentials (ERPs) in infants even 2- to 5-days after birth. Grossman et al., 2007 observed greater gamma power activation in 4-month-old infants in response to facial stimuli with direct gaze vs averted gaze, and where an experimenter engages in mutual gaze before looking towards an object, infants show enhanced neural processing of those objects (Parise et al., 2008; Hoehl et al., 2014). Of note, findings of gaze orientation on ERP amplitudes have not replicated well in developmental research; for example, Elsabbagh and colleagues (2009) found no significant effect of gaze type, comparing ERP amplitudes (see control group comparison); and findings are also largely mixed in adults (e.g., Watanbe et al., 2001; Taylor et al., 2001b; Watanbe et al., 2002; Itier et al., 2007; Conty et al., 2007; Ponkanen et al., 2011). Despite the inconsistencies, these findings have contributed to the popular idea that infants are highly sensitive to their partner’s social signals during early learning exchanges (Hains & Muir, 1996b; Symons et al., 1998; Farroni et al., 2004; Werker & Yueng, 2005; Werker et al., 2007; Hoehl et al., 2008; see Cetincelik et al., 2021 for a review). This raises basic questions of in what contexts, and under what circumstances are infants sensitive to their partner’s social signals.

One important limitation of research measuring infants’ neural responses to social stimuli presented on a screen, however, is its limited ecological validity. Historically, the majority of studies into early social development have measured infants’ passive responses to viewing a series of static images that flash on and off on a screen in a predetermined sequence. The real world, in contrast, is interactive, contingent and continuous. In recent years an increasing number of researchers have begun to recognise that, to study how the infant brain subserves social interaction, it is necessary to actually study it in interactive contexts (Schilbach et al., 2013; Redcay & Shilbach, 2019; Wass et al., 2020; Wass & Goupil, 2022). Recent behavioural findings have suggested important differences between screen-based simulacra of social interaction and actual social interaction. For example, recent studies (Franchak et al., 2011; Yu & Smith, 2013) have shown that infants rarely look to their caregiver’s face and eyes during free-flowing interactions, this is in contrast to what has previously been shown using screen-based tasks (e.g., Vecera & Johnson, 1995; Farroni et al., 2004). So far, the neural processing of mutual gaze has largely been investigated in un-ecological contexts, and in the absence of real social interaction. Consequently, two important questions remain unanswered; 1) is mutual gaze really a salient communicative signal during free-flowing social interactions occurring in rich, continuous, natural scenes? 2) how does intra and inter-brain activity support the processing of sender and receiver ostensive signals such as mutual gaze when both partners are engaged in a free-flowing, bidirectional exchange of information?

### Mutual gaze and inter-brain synchrony

Another topic that has shown rapidly burgeoning popularity in recent years is inter-brain synchrony. At the neural level, inter-brain synchrony can be defined as a dyadic mechanism, wherein temporally coordinated patterns of brain activity between two interacting individuals supports aspects of their ongoing social interaction (Holroyd 2022). A number of studies have observed increased inter-brain synchrony during mutual gaze. The majority of this research claims to measure inter-brain synchrony, although we recognise that not all of these studies will meet the framework of inter-brain synchrony set out in more recent theoretical accounts (Holroyd, 2022). Kinreich and colleagues (2017) observed significantly correlated gamma (30-60Hz) activity between interacting adults during social interaction. Higher interpersonal gamma correlations were also associated more strongly with mutual vs non-mutual gaze. Similarly, Luft and colleagues (2021) found that mutual gaze was associated with higher inter-brain gamma band (30-45Hz) coherence (a spectral measure based on correlation) between interacting adults than non-mutual gaze. In the developmental literature, our group investigated inter-brain synchrony in 7.5-month infant-adult dyads during moments of mutual and non-mutual gaze (Leong et al., 2017). During a live social, but not interactional condition infants observed an adult singing nursery rhymes, who was instructed to look either directly at the infant, directly at the infant with their head turned at a slight angle, or away from the infant. Consistent with research on adults, we found greater infant-adult neural synchrony during moments of mutual vs non-mutual gaze, measured using partially directed coherence (PDC-a spectral Granger causal measure of synchrony) in Theta (3-6Hz) and Alpha (6-9Hz) band activity. This study thus suggests that the impact of mutual gaze on inter-brain synchrony found in adult-adult dyads (Kinreich et al., 2017; Luft et al., 2021) is already present early on in development, though possibly in lower frequency brain rhythms.

### Sender/ Receiver mechanisms of inter-brain synchrony

As recent theoretical accounts have highlighted (Burgess 2013, Hamilton, 2021; Holroyd, 2022) inter-brain synchrony can reflect underlying mechanisms of varying complexity. To date, research investigating inter-brain synchrony during social interaction has exclusively measured this using non-event locked analyses, i.e., inter-brain synchrony values are averaged across whole conditions and/ or whole interactions and not time locked to any specific events within the interaction. Previously we have argued that in order to distinguish between different mechanisms that might give rise to inter-brain synchrony it is important to use event locked analyses (Haresign et al., 2022), i.e., analyses that focus on measuring fine-grained temporal changes in inter-brain synchrony, time-locked to specific behaviours/ events within social interactions. Additionally, when trying to differentiate inter-brain synchrony from other forms of inter-personal neural synchrony, it is also important to measure dyadic dynamics, e.g., how both partners’ behavior and neural activity contribute to establishing inter-brain synchrony.

We have suggested that one leading candidate mechanism for establishing inter-brain synchrony during mutual gaze may be mutual phase resetting (Wass et al, 2020; Leong et al., 2017). It is known that the phase of neuronal oscillations reflects the excitability of underlying neuronal populations to incoming sensory stimulation (Klimesch et al., 2007; Jensen et al., 2014). Consequently, there has been much effort expended in recent years, across a range of research fields, on exploring whether neuronal oscillations could be a key mechanism for temporal sampling of the environment (Schroeder & Lakatos, 2009; Giraud & Poepell 2012; Thut et al., 2012; VanRullen, 2016b; Ruzzoli et al., 2019). For example, some evidence suggests that sensory information arriving during high-receptivity periods is more likely to be perceived than information arriving during low-receptivity periods (Busch et al., 2009; Mathewson et al., 2009; 2010; 2011; 2012). This suggests that there is an optimal (range of) phase for perceiving information. It has been argued that if this is correct then it is logical to assume that there exist mechanisms, either endogenous or exogenous, for modulating the phase of neuronal oscillations, in order to match the temporal structure of the environmental input (van Diepen et al., 2015; Ruzzoli et al., 2019). This mechanism (phase resetting) would enable more efficient processing as information would be received during periods (phases) of high receptivity.

Empirical evidence supports the role of phase resetting as an intra-brain mechanism, facilitating neural entrainment to temporal structures within the environment, for example speech (Giraud & Poepell 2012; Biau et al., 2015). It is therefore logical to question whether similar mechanisms also operate at the interpersonal level. For example, inter-brain synchronisation may increase within a dyad following the onset of communicative signals (such as gaze, gestures, or vocalisations) that reset the phase of both interacting partners. Here, neural oscillations in both the sender (of the social signal) and the receiver’s brain that were previously random with respect to each other (low inter-brain synchrony) would be simultaneously reset in response to a communicative signal. Following this reset the neural activity of both the sender and the receiver would oscillate with more consistent variation over time (high inter-brain synchrony). We recognise that according to recent theoretical accounts that this might be classed as motor-induced neural synchrony, which occurs when the behaviour of one member of the dyad drives the neural activity of both members of the dyad (Holroyd, 2022). However, it could also be that mutual gaze onsets reset the brain activity of the sender, which precedes/ causes the behaviour of the receiver, which then causes a reset in the receiver’s brain activity. Here, increases in inter-brain synchrony would be a result of both partners resetting to their own behaviours. Distinguishing between these different mechanisms is only possible using event locked analysis.

### Eye movement (saccade) related potentials and ERPs

One challenge in studying the impact of mutual gaze during naturalistic social interaction on dyadic EEG is that mutual gaze onsets are time-locked to eye movements which create multiple types of artifact in the EEG (Dimigen, 2020). For example, PlÖchl and colleagues (2012) showed that saccadic spike potentials (EEG potentials time-locked to small, < 1°, involuntary eye movements during fixation) typically introduce a broadband artifact in the time-frequency spectrum of the EEG, which is strongest (amplitude) in the low beta (∼14-30 Hz) and gamma bands (>30 Hz) of adult EEG and typically peaks between -50ms and 150ms around the offset of a saccade. Artifact generated from eye movements can overlap in time and frequency with EEG activity presumed to be related to genuine neural activity, associated with stimulation of the retina (Gaarder et al., 1964; Billings, 1989; Dimigen, 2021). This is often referred to as the lambda response (LR) (Kazai & Yagi, 2003), which is an occipital EEG potential that can be observed when saccades are made against an illuminated contrast background (Thickbroom et al., 1991). LRs typically produce broadband time-frequency activity that is strongest (amplitude) in Alpha (8-13 Hz) and low beta (∼14-30 Hz), over occipital electrodes and peaks ∼100ms after the offset of the saccade (Dimigen et al., 2009; 2011). The overlapping activations introduced by eye movements can make interpretation of the data challenging, a problem which is not solved using ‘standard’ artifact correction procedures which fail to completely remove artifact associated with eye moments from the EEG; both in adults (PlÖchl et al., 2012; Dimigen, 2020) and infants (Haresign et al., 2021).

Our analyses are presented using a pipeline specially designed for the removal of eye movement artifact from naturalistic EEG data using ICA (Georgieva et al., 2020; Haresign et al., 2021; Kayhan et al., 2022). However, it is important to note that in our 2021 paper we reported that we (as arguably most of the current research in developmental neuroscience using EEG is) were unable to completely remove the activity that we assumed to be artifactually related to eye movements. Therefore, in this current work, it is likely that the sender neural responses that we are investigating are a combination of some residual artifactual activity; although as discussed above these artifacts are transient (∼100ms) and therefore would only impact the initial part of the ERP waveform, and genuine neural activity; after the initial ∼+150-200ms. For this reason, our primary analyses will compare sections of our data that both contain saccades, and therefore have (we assume) an identical amount of eye movement artifact in them but have different consequences (either the saccade leads to mutual gaze, or not). Furthermore, we only compared activity in the later parts of the ERP waveform after the first 100ms.

### Current study: The role of mutual gaze onsets in creating inter-brain synchrony

Our study aimed to test the hypothesis that infants are sensitive to ostensive signals during free-flowing social interactions, and that mutual gaze onsets lead to mutual phase resetting, which causes increases in inter-brain synchrony. We measured dual EEG recordings from parents and infants whilst they engaged in free-flowing social interactions and investigated intra- and inter- individual neural responses to naturally occurring moments of mutual gaze. To explore the role of turn-taking in creating inter-brain synchrony, we differentiated between two types of look onset, depending on each partners’ role. ‘Sender’ gaze onsets were defined at a time when either the adult or the infant made a gaze shift towards their partner at a time when their partner was either already looking at them (sender mutual) or not looking at them (sender non-mutual). ‘Receiver’ gaze onsets were defined at a time when their partner made a gaze shift towards them at a time when either the adult or the infant was already looking at their partner (receiver mutual) or not (receiver non-mutual) (see Figure 1).

**Figure 1.**
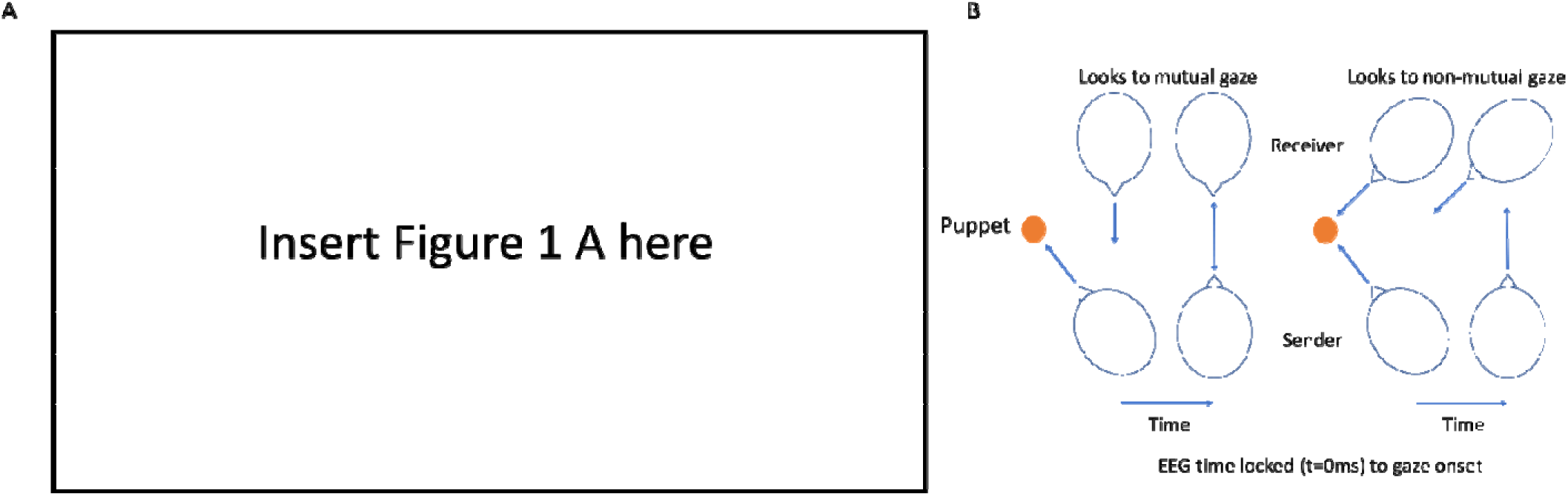
Illustration of experimental set-up and design for event locked analysis. A) shows screenshots of experimental recordings from three camera angles used. B) shows the design of the event locked analysis. Sender gaze onsets were defined at a time when either the adult or the infant made a gaze shift towards their partner’s face at a time when their partner was either reciprocating their gaze (sender mutual gaze onsets) or not reciprocating their gaze (sender non-mutual gaze onsets). Receiver gaze onsets were defined at a time when either the adult or the infant was already looking at their partner’s face (receiver mutual gaze onsets) or not looking at their partner (receiver non-mutual gaze onsets and their partner made a gaze shift towards them. During receiver non-mutual gaze onsets, the receiver was looking at the puppet.

In order to assess how these interpersonal dynamics contributed to inter-brain synchrony we used two different measures of synchrony; First, we used a measure of concurrent synchrony, Phase locking value (PLV), that measures zero-lag, undirected synchrony. This measure would best capture changes in inter-brain synchrony that resulted from changes in both partners’ brains concurrently. We also used a measure of sequential synchrony, Partially directed coherence (PDC), that measures time lagged, directed synchrony. This would best capture changes in inter-brain synchrony that resulted from changes in one partner’s brain that forward predicted or lead to changes in the other partner’s brain. Through this we were able to consider three main sets of research questions.

### Inter-brain non-event locked analysis

For our first set of research questions, we take an analytical approach similar to Leong and colleagues’ 2017 paper, in which we explore whether inter-brain synchrony is stronger overall during moments of mutual gaze. Although this was not a replication study, we attempted to translate previous structured experimental designs into a more naturalistic context. For these analyses, we preselected frequency bands and electrodes of interest, based on the findings of Leong and colleagues (2017). Consistent with these findings we expected to observe greater inter-brain synchrony, in Theta and Alpha during all moments of mutual vs non-mutual gaze. Inter-brain synchrony was measured using PLV and PDC computed over EEG data, averaged over central electrodes (C3 and C4).

### Inter-brain event-locked analysis

Investigating inter-brain synchrony during social interactions as a time invariant phenomenon makes it difficult to understand the underlying mechanisms (Haresign et al., 2022). Therefore, for our second set of research questions, we wanted to explore how inter-brain synchrony changes around gaze onsets. We compared inter-brain synchrony values around sender and receiver mutual vs non-mutual gaze onsets. We expected to observe greater inter-brain synchrony (measured using PLV and PDC, in frequencies 2-18 Hz, over occipital electrodes) around mutual vs non-mutual gaze onsets. For all inter-brain analyses, we used one measure of concurrent (PLV) and one measure of sequential (PDC) synchrony in order to try to better understand how sender and receiver dynamics influence inter-brain synchrony.

### Intra-brain event locked analysis

It has been suggested that one mechanism that might mediate changes in inter-brain synchrony is mutual phase resetting in response to the onset of mutual gaze (Leong et al., 2017, Wass et al., 2020). For our third set of research questions, we wanted to further investigate, mechanistically how changes in inter-brain synchrony might develop around mutual gaze onsets. To examine this, we first looked at ERPs and inter-trial coherence (ITC) around gaze onsets: comparing sender and receiver mutual and non-mutual gaze onsets. Inter-trial coherence measures consistency in phase angles over trials/ time at a single given electrode and has been used to phase resetting (Makeig et al., 2002). In comparison PLV measures the consistency in phase angle differences between two electrodes. Based on previous findings (e.g., Farroni et al., 2002) we expected to observe significant ERPs and increases in ITC relative to both sender and receiver mutual and non-mutual gaze onsets. Although during receiver non-mutual gaze onsets only, the receiver was looking at an object and not at their partner’s face, research has shown that humans are highly sensitive to eye gaze in their peripheral vision (Loomis et al., 2008). ERPs and ITC were first assessed against a baseline to test whether there was a significant event-locked neural response relative to both sender and receiver mutual and non-mutual gaze onsets. We then examined whether ERPs and event locked ITC was greater around sender and receiver mutual vs non-mutual gaze onsets. We expected to observe larger ERP amplitudes, and greater ITC (in frequencies 2-18 Hz, over occipital electrodes) around mutual vs non-mutual gaze onsets.

## 2. Methods

### 2.1. Ethics statement

This study was approved by the Psychology Research Ethics Committee at the University of East London. Participants were given a £50 shopping voucher for taking part in the project.

### 2.2. Participants

Of the 90 infants we tested for this study, 21 contributed no data at all, 6 contributed EEG data that was too noisy even after data cleaning and 4 were lost due to human error, e.g., failed synchronisation triggers. We also excluded all participants with fewer than 5 gaze onsets, leading to an additional 4 datasets being excluded. The final sample contained 55 healthy (23 F), *M* = 12.2-month-old (*SD =*1.47) infants, that participated in the study along with their mothers.

### 2.3. Power calculations

For the non-event locked analysis, as this analysis were based on previous findings, we estimated the required sample size to observe a difference between the two groups (as a product of gaze type), using the G*power tool (Faul et al., 2007). For this, we used data from Leong et al., (2017) as an estimator of the expected effect size for the analysis of non-event locked synchrony (0.332). Based on an Alpha level of .05, in our sample size (*N* = 55) we had a >99% chance of observing an effect of gaze type on inter-brain synchrony of the magnitude observed in previous work.

### 2.4. Experimental set-up and procedure

Infants were positioned immediately in front of a table in a highchair. Adults were positioned on the opposite side of the 65cm-wide table, facing the infant. Adults were asked to stage a ‘three-way conversation’ between the infant and a small hand puppet and to try to spend an equal amount of time looking at the puppet and the infant. Dual EEG was continuously acquired from the parents and infants for the approx. 5 min duration of the play session (M = 386.1, SD = 123.9 seconds)

### 2.5. Behavioural data

Video recordings were made using Canon LEGRIA HF R806 camcorders recording at 50fps positioned next to the infant and parent respectively. Video recordings of the play sessions were coded offline, frame by frame, at 50 fps. This equates to a maximum temporal accuracy of ∼20ms. Coding of the infant’s and adult’s gaze was performed by two independent coders. Cohen’s kappa between coders was >85%, which is high (McHugh, 2012). EEG was time-locked to the behavioural data offline based on the video coding using synchronized LED and TTL pulses. To verify the synchronisation, we manually identified blinks in the behavioral data and looked to see if this matched the timing of the blinks in the EEG data.

### 2.6. EEG data acquisition

EEG signals were obtained using a dual 32-channel Biosemi system (10-20 standard layout), recorded at 512 Hz with no online filtering using the Actiview software.

### 2.7. EEG artifact rejection and pre-processing

A fully automatic artifact rejection procedure was adopted, following procedures from commonly used toolboxes for EEG pre-processing in adults (Mullen, 2012; Bigdely-Shamlo, et al., 2015) and infants (Gabard-Durham et al., 2018; Debnath et al., 2020). Full details of the pre-processing procedures can be found in (Haresign et al., 2021). In brief the data was filtered between 1 and 20Hz and re-referenced to a robust average reference. Then we interpolated noisy channels based on correlation; if a channel had a lower than 0.7 correlation to its robust estimate (average of other channels) then it was removed. The mean number of channels interpolated was 3.9 (*SD =*2.1) for infants and 3.9 (*SD =*4.4) for adults. Then for the infant data only we removed sections from the continuous data in which the majority of channels contained extremely high-power values. Data was rejected in a sliding 1 second epoch and based on the percentage of channels (set here at 70% of channels) that exceeded 5 standard deviations of the mean EEG power over all channels. For example, if more than 70% of channels in each 1-sec epoch exceeded 5 times the standard deviation of the mean power for all channels then this epoch is marked for rejection. We found that for adults this step was primarily removing activity that could be removed with ICA (e.g., eye movement artifact) without removing entire sections of the data. The average amount of continuous data removed was 11.9% (*SD =*14.6%) for infants. Finally, we used ICA to remove additional artifacts.

Careful attention was paid to artifact and the amount of noise in the data throughout. In the supplementary materials (see SM 8) we report the results of standard measures of EEG data quality (Luck et al., 2021). Including universal measures like these enables fast and easy comparison between studies and allows the overall quality of the data to be readily assessed.

We paid particular attention to eye movement artifact. In previous work we designed a system for automatically identifying and removing artifactual ICA components in infant EEG (Haresign et al., 2021). The automated system was shown to remove most but not all eye movement related artifact time-locked to saccades. Therefore, we cannot entirely rule out that some of the activity in the sender brains relative to sender mutual and non-mutual gaze onsets is an artifact of the gaze shift.

### 2.8. Time frequency analysis- extracting power and phase

Time-frequency power and phase was extracted via complex Morlet wavelet convolution. The wavelets increased from 2 to 18 Hz in 17 linearly spaced steps and the number of cycles increased from 3-10 cycles logarithmically (this approach is generally recommended; see Cohen, 2014, chapter 13).

For all non-event locked analyses presented here, frequency bands were selected based on the bands commonly used in infant research: Theta (3-6Hz) and Alpha (6-9Hz) (Marshall et al., 2011; Leong et al., 2017; Xie & Richards, 2018; van der Velde et al., 2019; Jones et al., 2020).

## 3. Analysis

### 3.1. Inter-brain non-event locked analysis - overview

First, we wanted to explore whether inter-brain synchrony was greater during all moments of mutual vs non-mutual gaze (not relative to sender/ receiver gaze onsets). Mutual gaze was defined as times when both the adult and the infant were looking at each other. Non-mutual gaze was defined as times when the infant was looking at the adult and the adult was not looking at the infant (or vice versa). For all non-event locked analyses, EEG was time-locked to the onset of gaze, and the length of the epoch extracted equated to the duration of the look. The average look durations were 2.7 seconds (SD = 0.9s) for mutual and 1.6 seconds (SD = 0.5s) for non-mutual gaze. All epochs were then concatenated. The mean amount of continuous data available for analysis was 66.1s (SD = 41.5s) for mutual and 28.3s (SD = 17.9s) for non-mutual gaze. Because ITC (see Cohen 2014, chapter 19) and PDC are sensitive to the amount of available data, we normalised the amount of data present per condition for each participant by identifying which gaze type had the lower number of continuous data samples (n), and re-sampling data from the other gaze type condition, taking 1:n data samples.

#### 3.1.1. Inter-brain non-event locked analysis - PDC

Partial directed coherence (PDC) is based on the principles of Granger causality (Baccalá and Sameshima, 2001). It provides information of the extent to which one times series influences another. PDC is calculated from coefficients of autoregressive modelling according to:

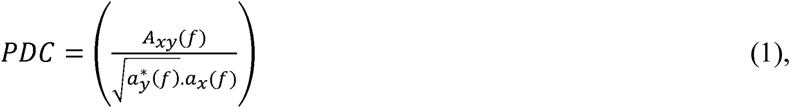

where *A_xy_(f)* is the spectral representation of bivariate model coefficients and *a_y_* and *a_x_* are the spectral model coefficient from the univariate autoregressive model fit. Based on previous literature (e.g., Leong et al., 2017) we chose to compute PDC in 1-second non-overlapping sliding window. We estimated the model order for each segment using Bayesian information criteria (BIC). Model order values were then averaged for all segments. The result was a model order of 5, the same as used by Leong and colleagues (2017), which was then used for all segments.

#### 3.1.2. Inter-brain non-event locked analysis – PLV

For the non-event locked analyses, the phase locking value (PLV) was calculated within a single trial over a defined temporal window (e.g., Tass et al., 1998) according to:

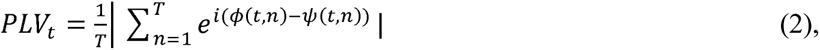

Where T is the number of observations or time samples within the window, *ϕ*(*t,n*) is the phase on observation *n*, at time *t*, in channel *ϕ* and *ψ(t,n)* at channel *ψ*.

#### 3.1.3. Inter-brain non-event locked analysis – group level stats for PLV and PDC

Firstly, we assessed whether PLV and PDC values were significant compared to a baseline level. This was done by calculating the PLV/ PDC between randomly paired infant and adult dyads. We generated 1000 random infant-adult pairings in this way. We then compared the group averages of the observed PLV/ PDC values for the real infant-adult pairings against the randomly permuted distributions. Under the null hypothesis that the interbrain PLV/ PDC between infants and adults is a result of their real-time social interaction, we should observe no above chance inter-brain PLV/ PDC between randomly paired infant-adult dyads. P values were generated by first z-scoring the observed PLV values and then by evaluating the z-scored value’s position on a Gaussian probability density using the Matlab function normcdf.m. For this test, we used a Bonferroni correction for multiple comparisons. This was appropriate here as we were only testing over a limited number of predefined frequencies/ channels. Following the statistical procedure adopted by Leong and colleagues’ (2017), differences in PLV and PDC between mutual and non-mutual gaze were assessed using a two-way repeated-measures ANOVA, taking gaze type and frequency as the within levels, using average, over electrodes C3 and C4, infant-to-adult PDC *(I*➔*A)* and adult-to-infant *(A*➔*I)* PDC values, and for Theta and Alpha bands separately. A Tukey-Kramer correction for multiple comparisons was applied.

### 3.2. Inter-brain event locked analysis

#### 3.2.1. Inter-brain event locked analysis – group level stats for PLV and PDC

Firstly, we assessed whether PLV and PDC values were significant compared to a baseline level. We compared all observed time-frequency PLV/ PDC values relative to sender and receiver mutual and non-mutual gaze onsets against time-frequency PLV/PDC values time-locked to randomly inserted events within the continuous data. Differences between the real and surrogate data were assessed using cluster-based permutations statistics, using an alpha value of .025 (see SM section 5 for full details). Secondly, we examined whether PLV/ PDC values were greater for sender/ receiver mutual vs non-mutual gaze onsets. This was similarly assessed over all time-frequency points using a cluster-based permutation procedure, comparing results between different types of gaze onset. Importantly, the amount of eye movement artifact was identical between the sets of results being compared.

### 3.3. Intra-brain event locked analysis

Here we wanted to investigate whether mutual gaze onsets play a role in establishing inter-brain synchrony. We did this by investigating ERPs and ITC around sender and receiver mutual and non-mutual gaze onsets. Previous research suggests that event-locked face-sensitive neural responses are strongest over parietal/ occipital electrodes (Gao et al., 2019; Haresign et a., 2021). Therefore, for our event locked analysis we chose to focus on averaged data from a cluster of 5 parietal/ occipital electrodes (PO3, PO4, O1, Oz, O2). Additional topoplots are presented in SM 7, which support the choice of electrodes. The EEG signal was divided into events from -2500 to 2500ms (t=0 denotes the onset of gaze). The mean number of events extracted was 21.1 (SD =10.8) for infant sender/ adult receiver and 15.6 (SD =12.6) for adult sender/ infant receiver mutual gaze onsets and 9.9 (SD =6.2) for infant sender/adult receiver and 28 (SD =18.3) for adult sender/infant receiver non-mutual gaze onsets. We matched the number of events between gaze types for each participant using the procedure described in section 3.1, above.

#### 3.3.1. Intra-brain event locked analysis - Inter trial coherence

Inter trial coherence (ITC) measures the consistency of frequency band-specific phase angles over trials, time-locked to a specific event. The phase coherence value is computed according to:

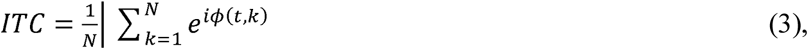

where *N* is the number of trials and *ϕ*(*t,k*) is the phase on trial *k*, at time *t*.

#### 3.3.2. Intra-brain event locked analysis - ERPs

Following previous research (Conte et al., 2020) amplitudes of the P1, N290, and P400 ERPs were measured by calculating the change in amplitude between the peak of the component of interest and the peak of the preceding component. Also following previous recent research (Conte et a., 2020; Guy et al., 2016; 2018; Xie and Richards, 2016; 2017) we used semiautomated and individualised time window selection (Guy et al., 2021). Differences in peak amplitude were quantified using the adaptive mean approach. This process involves first identifying the peak latency of the ERP potential on a participant-by-participant basis using a broad time window. Once the peak latency has been identified an average of the activity in a 20ms window around the peak is then taken (e.g., as described in Luck, 2014). We focused on three major components relevant for face/gaze processing: the P1 component, the N170 (commonly N290 in infant EEG; Conte et al., 2020) and the P300 (commonly P400 in infant EEG; Conte et al., 2020). For the P1 component we used a time window of 0 to 200ms for adults and 100 to 300ms for infants. For the N170/ N290 component we used a time window of 100 to 300ms for adults and 200 to 400ms for infants. For the P300/P400 component we used a time window of 200 to 500ms for adults and 300 to 600ms for infants. These were selected based on visual inspection of the averaged waveforms. All ERP data were baseline corrected using data from the time window -1000 to -700ms pre-gaze onset.

#### 3.3.3 Intra-brain event locked analysis – group level stats for ERPs

To test whether the onset of gaze led to significant changes in amplitude relative to both sender and receiver mutual and non-mutual gaze onsets we again used nonparametric permutation testing. Here the null hypothesis was that the timing of the gaze onset (e.g., time 0) is unrelated to the observed neural response within the time window examined. To test this, we randomly permuted the time points of the ERPs and took the average (separately around the maximum and minimum points) of the permuted ERP in the time window 0 to +500ms. This procedure was then repeated 1000 times, randomising and reshuffling the ERP on each permutation. Finally, an estimate of the permutation p-value was calculated using the z-scoring procedure outlined in section 2.12. Here we used cluster-based permutation statistics, using an Alpha value of .025 to correct for multiple comparisons. The results are reported in section 4.3.

#### 3.3.4 Intra-brain event locked analysis – group level stats for ITC

Firstly, we assessed whether ITC values were significant compared to a baseline level. We compared the observed time-frequency ITC values relative to sender and receiver mutual and non-mutual gaze onsets against time-frequency ITC values time-locked to randomly inserted events within the continuous data. The number of random events was matched based on the number of real gaze onsets available for each participant. Differences between the real and surrogate data were assessed using cluster-based permutation statistics, using an Alpha value of .025 (see SM 4 for full details). We then compared differences between sender and receiver mutual vs non-mutual gaze in ITC using the same cluster-based permutation procedure see section 4.4.

## 4. Results

Before turning to our main research questions, we first calculated descriptive statistics to show how gaze onsets were distributed in our sample (Figure 2). These results indicate that infants spent, in total, 34%/31%/35% of the total interaction looking to partner/puppet/inattentive (Fig 2A). Parents spent 62%/31%/7% of the total interaction looking to partner/puppet/inattentive (Fig 2D). Overall, mutual gaze periods were longer than non-mutual gaze periods (Fig 2C). Differences in the frequency of gaze onsets to different types of gaze were normalised using the procedures described above.

**Figure 2.**
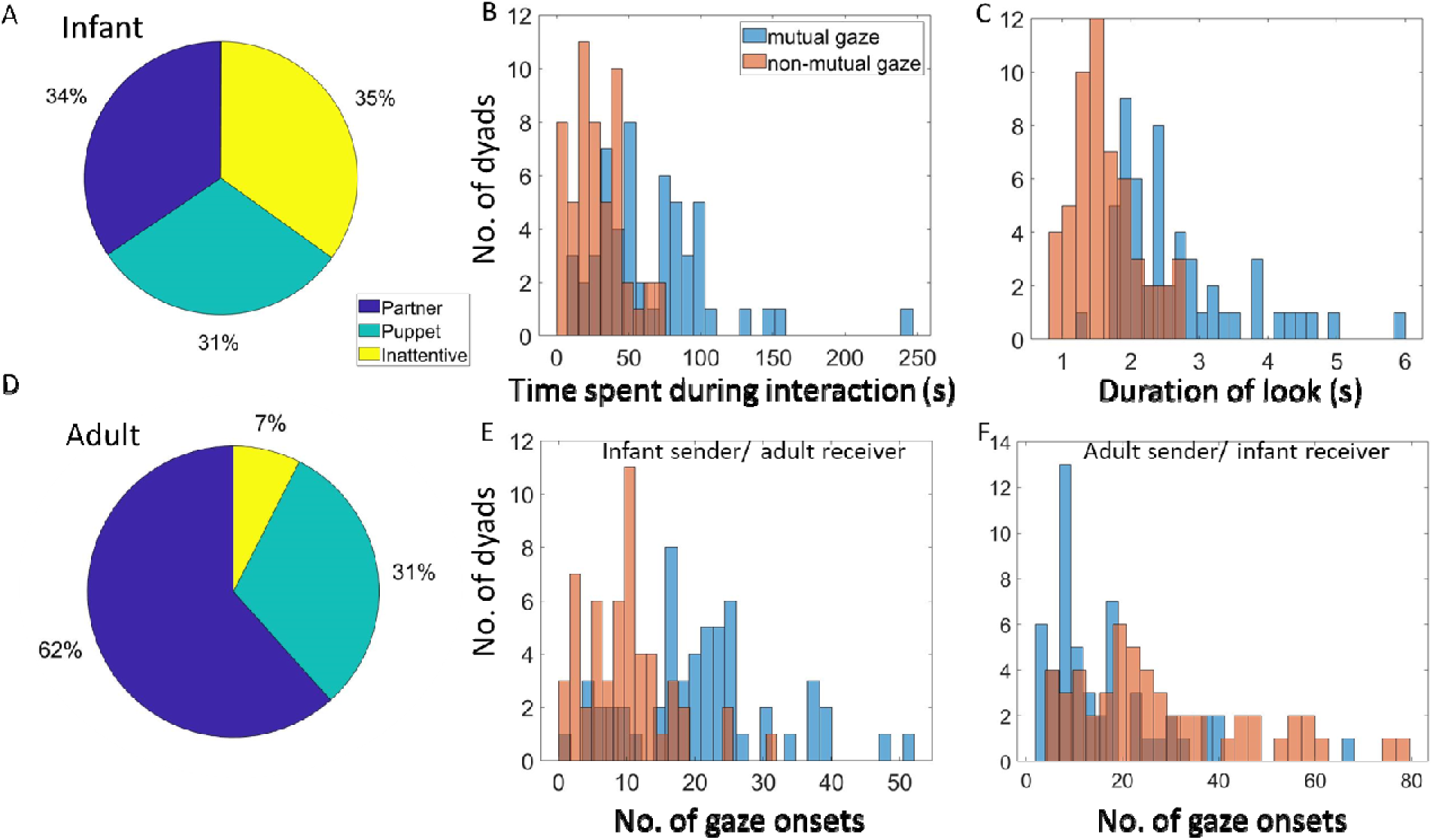
Distribution of gaze onsets in our sample. A) Distribution of time infants spent looking at different areas. B) Distribution of time dyads spend in mutual and non-mutual gaze during interaction. C) Distribution of look durations for mutual and non-mutual gaze defined from infants’ look behaviour. D) Distribution of time adults spent looking at different areas. E) Distribution of infant sender/ adult receiver mutual and non-mutual gaze onsets. F) Distribution of adult sender/ infant receiver mutual and non-mutual gaze onsets.

### 4.1. Inter-brain non-event locked analysis – PLV and PDC

To investigate the relationship between inter-brain synchrony and mutual gaze, we first computed the mean PLV and PDC values across all mutual and non-mutual gaze periods in Theta and Alpha bands separately. We looked at whether PLV and PDC values were significantly greater than baseline, and then compared these values for mutual vs non-mutual gaze. Based on our previous research (Leong et al., 2017) we focused on activity over vertex electrodes (C3 and C4).

Figures 3 and 4 show the results of the inter-brain non-event locked analysis. We first tested whether PLV and PDC values significantly exceeded baseline values. The results of the permutation analysis indicated that PLV values did not significantly exceed baseline values: in theta (3-6 Hz) for mutual (*p* = 0.53) or non-mutual gaze (*p* = 0.56). Further, infant-to-adult PDC *(Infant*➔*Adult)* and adult-to-infant *(Adult*➔*Infant)* did not significantly exceed baseline values for mutual (*p* = 0.63/ *p* = 0.55) or non-mutual gaze (*p* = 0.64/ *p* = 0.54); or in alpha (6- 9 Hz) for mutual (*p* = 0.57) or non-mutual gaze (*p* = 0.59). Further, infant-to-adult PDC *(Infant*➔*Adult)* and adult-to-infant *(Adult*➔*Infant)* did not significantly exceed baseline values for mutual (*p* = 0.59/ *p* = 0.53) or non-mutual gaze (*p* = 0.61/ *p* = 0.52).

**Figure 3.**
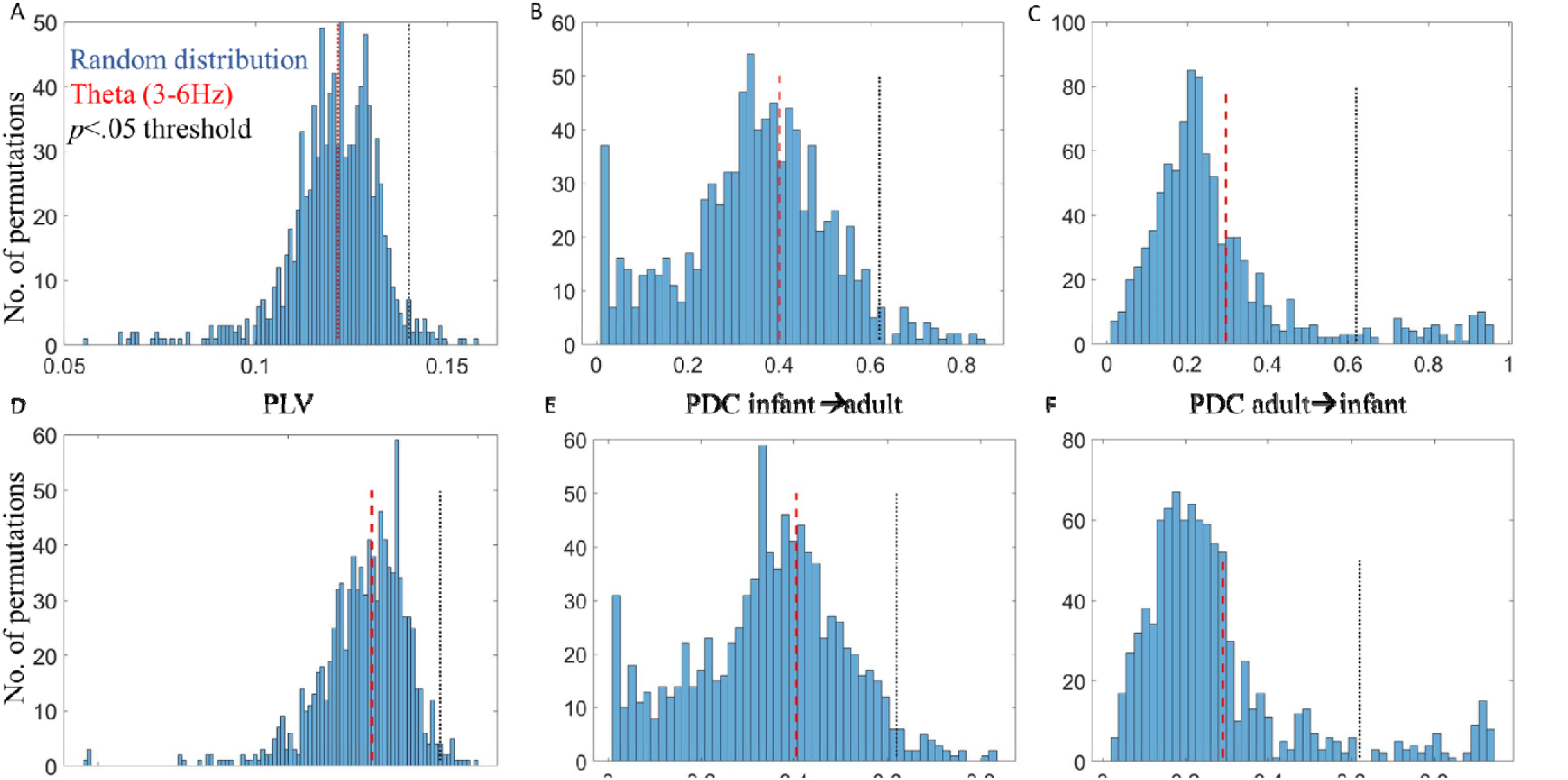
Results of permutation tests for non-event locked inter-brain synchrony analysis. A) Distribution of random pair infant-adult PLV values compared to real pair infant-adult PLV values in Theta for mutual gaze. B) Distribution of random pair infant→adult PDC values compared to real pair infant→adult PDC values in Theta for mutual gaze. C) Distribution of random pair adult→infant PDC values compared to real pair adult→infant PDC values in Theta for mutual gaze. D) Distribution of random pair infant-adult PLV values compared to real pair infant-adult PLV values in Theta for non-mutual gaze. E) Distribution of random pair infant→adult PDC values compared to real pair infant→adult PDC values in Theta for non-mutual gaze. F) Distribution of random pair infant→adult PDC values compared to real pair adult→infant PDC values in Theta for non-mutual gaze. Red dashed lines indicate averaged observed values in Theta, black dotted lines indicate the threshold for p < 0.05.

**Figure 4.**
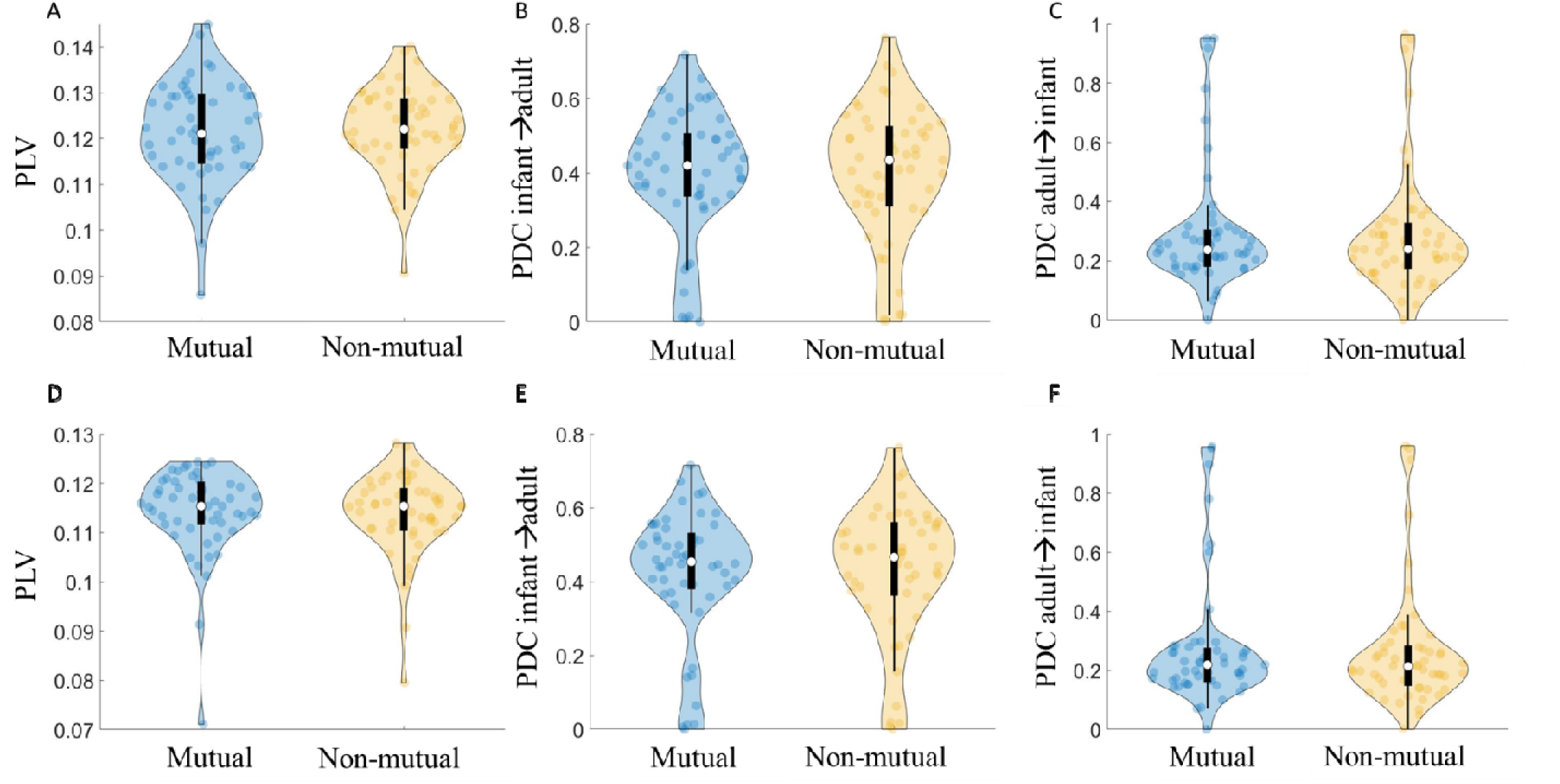
Results of non-event locked inter-brain synchrony analysis. A) Infant-adult PLV during mutual / non-mutual gaze in Theta. B) Infant→adult PDC during mutual / non-mutual gaze in Theta. C) Adult→infant PDC during mutual / non-mutual gaze in Theta. D) Infant-adult PLV during mutual / non-mutual gaze in Alpha. E) Infant-adult PLV during mutual / non-mutual gaze in Alpha. F) Adult→infant PDC during mutual / non-mutual gaze in Alpha. Violin plots show the distribution of the data with an inset boxplot. Each point corresponds to the average PLV/PDC value of a dyad.

We then wanted to test whether PLV and PDC values were greater for mutual than non-mutual gaze. As described in section 3.1.3, we then conducted repeated measures (RM) ANOVAs using average indices (over C3 and C4), taking frequency (2 levels) and gaze type (mutual vs non-mutual; 2 levels) as within-subjects factors. The results of the ANOVA indicated no statistically significant differences in PLV, *F*(1, 55) = .04, *p* = .84, or PDC; for infant to adult (*Infant*➔*Adult*) influences, *F*(1, 55) = .18, *p* = .68 or adult to infant (*Adult*➔*Infant*) influences, *F*(1, 55) = .50, *p* = .48, between mutual and non-mutual gaze. The results did indicate a significant effect of frequency (i.e., more synchrony in Theta than Alpha) for PLV, *F*(1, 55) = 47.33, *p* < .01 and PDC; *Infant*➔*Adult*, *F*(1, 55) = 41.2, *p* <.01 and *Adult*➔*Infant*, *F*(1, 55) = 138.84, *p* < .01). These results are summarised in Figure 4. Note these are uncorrected p values.

To further test the significance of gaze type on non-event locked synchrony (PLV and PDC) PDC) including both directions of influence. We calculated *BF*_10_ = *p*(*D|H*_1_)/*p*(*D*|*H*_0_), we calculated the Bayes Factor (BF) at the group level for both Theta and Alpha and (for where *D* represents the data and *H*_1_ and *H*_0_ of the alternative and the null hypothesis respectively, using functionality from the bayesFactor toolbox (Krekelberg, 2022), based on the equations provided in Rouder et al., 2012. The BF10 tests for the presence of an effect.

For all tests, the BF10 was between 1/3 and 1/10 and non-significant indicating moderate evidence (Lee & Wagenmakers, 2014) for the null hypothesis (that there is no difference between mutual and non-mutual gaze). We also converted these scores to the Bayes Factor for the absence of an effect (BF01), confirming that there was moderate to strong evidence that there was no difference between the groups. Results of our Bayes Factor analyses are given in full in SM section 6.

To summarise the results of the non-event locked analyses, suggest that mutual gaze does not induce inter-brain synchrony in this dataset and using this paradigm since a) the inter-brain synchrony for either gaze type isn’t above shuffled baseline data and b) isn’t different from a specifically selected other condition (non-mutual gaze).

### 4.2. Inter-brain event locked analysis – PLV and PDC

We next investigated whether onsets of mutual gaze led to changes in inter-brain synchrony and examined the sender-receiver dynamics that might contribute to this. To do this we conducted event-locked analyses with respect to gaze onsets. We first examined whether PLV and PDC values were significantly greater than baseline around gaze onsets, and then compared these values between mutual vs non-mutual gaze onsets.

We first tested whether PLV and PDC values over occipital electrodes and in the 2-18 Hz range significantly exceeded baseline values generated from a permutation procedure for infant sender/ adult receiver and adult sender/ infant receiver looks to mutual and non-mutual gaze. The result of the permutation analysis indicated that event locked PLV and PDC values around mutual and non-mutual gaze onsets were not significantly different from baseline values (see SM section 5 for full details). Therefore, we failed to reject the null hypothesis that there are no changes in PLV and PDC that are time-locked to gaze onsets.

We also observed no statistically significant differences for the effect of gaze type (e.g., mutual vs non-mutual). Therefore, we failed to reject the null hypothesis that there was no difference in PLV and PDC between looks to mutual or looks to non-mutual gaze (see Figure 4, and SM section 5).

To further test the significance of gaze type on event locked inter-brain synchrony (PLV and PDC) we calculated the Bayes Factor at the group level for both Theta and Alpha and (for PDC) including both directions of influence. For all tests, the BF10 was between 1/3 and 1/10 and non-significant indicating moderate evidence (Lee and Wagenmakers 2014) for the null hypothesis (that there is no difference between mutual and non-mutual gaze). We also converted these scores to the Bayes Factor for the absence of an effect (BF01), confirming that there was moderate to strong evidence that there was no difference between the groups. Results of our Bayes Factor analyses are given in full in SM section 6.

### 4.3. Intra-brain analysis - ERPs

Earlier we discussed some of the potential mechanisms that could lead to changes in inter-brain synchrony. Here, we wanted to examine intra-brain sender/ receiver dynamics around mutual gaze. To do this we compared ERPs between sender and receiver mutual vs non-mutual gaze onsets.

Figure 6 shows the results of the intra-brain ERP analysis, comparing sender and receiver mutual and non-mutual gaze onsets. We first tested whether ERP values for sender and receiver mutual and non-mutual gaze onsets significantly exceeded baseline values generated from a permutation procedure (see section 3.3.3). The permutation analysis indicated that ERP amplitudes (this is just looking at whether there is a positive peak in the 0-500ms time window) in the post gaze onset window were significantly higher than baseline for sender mutual (*p* < 0.01 for both infants and adults) and non-mutual (*p* < 0.01 for both infants and adults) gaze onsets, in both infants and parents, but not for receiver mutual (*p = P_z_,* for infants and *p =* 0.2, for adults) or non-mutual (*p = P_z_,* for infants and *p =* 0.6, for adults) gaze onsets in either parents or infants.

**Figure 5.**
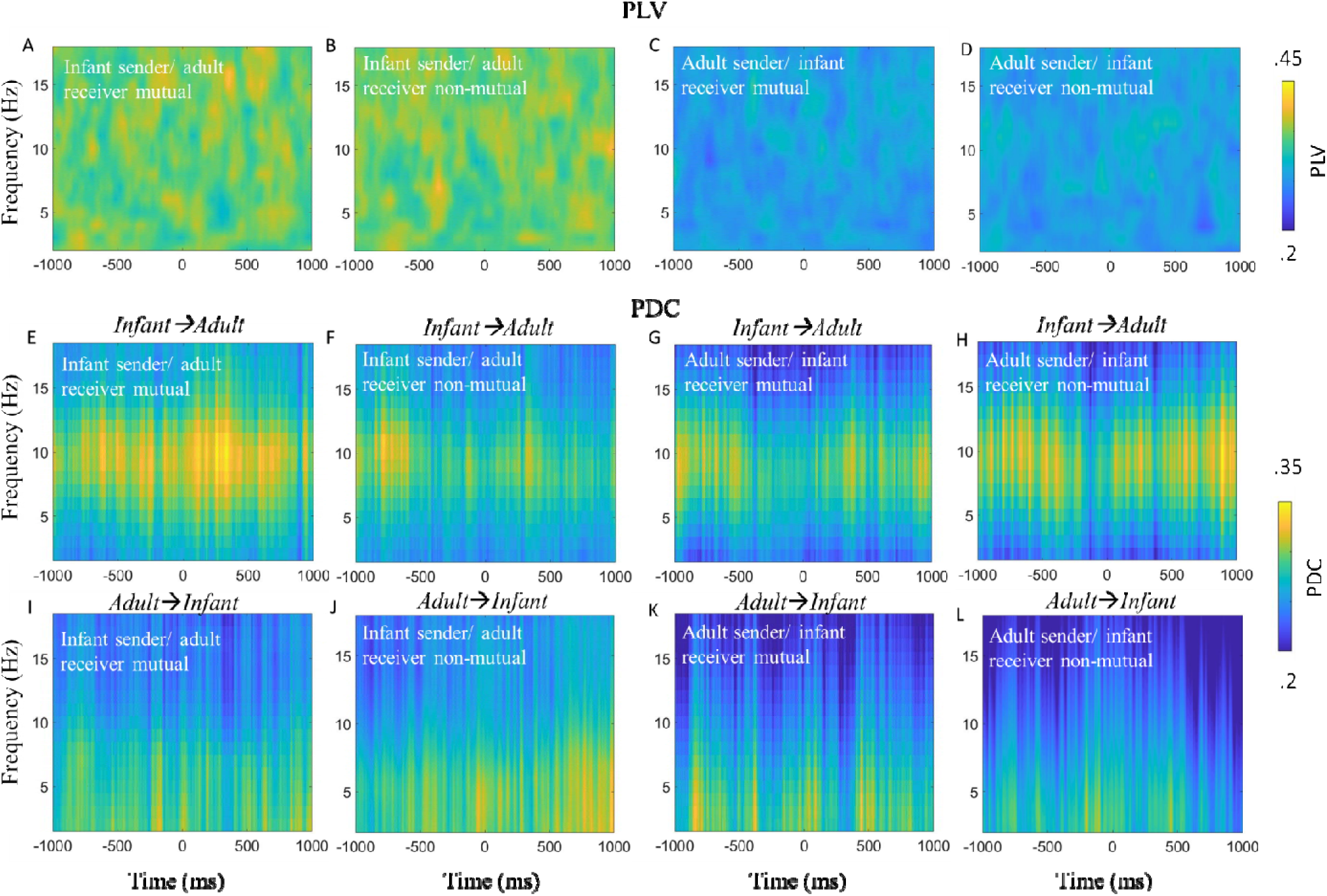
Infant-caregiver inter-brain synchrony time-locked to naturally occurring mutual and non-mutual gaze onsets. A) PLV relative to infant sender/adult receiver mutual gaze onsets. B) PLV relative to onsets of infant sender/adult receiver looks to non-mutual gaze. C) PLV relative to adult sender/infant receiver mutual gaze onsets. D) PLV relative to adult sender/infant receiver non-mutual gaze onsets. E) Infant→Adult PDC relative to infant sender/adult receiver mutual gaze onsets. F) Infant→Adult PDC relative to infant sender/adult receiver non-mutual gaze onsets. G) Infant→Adult PDC relative to adult sender/infant receiver mutual gaze onsets. H) Infant→Adult PDC relative to adult sender/infant receiver non-mutual gaze onsets. I) Adult→Infant PDC relative to infant sender/adult receiver mutual gaze onsets. J) Adult→Infant PDC relative to infant sender/adult receiver non-mutual gaze onsets. K) Adult➔Infant PDC relative to adult sender/infant receiver mutual gaze onsets. L) Adult➔Infant PDC relative to adult sender/infant receiver non-mutual gaze onsets. No significant differences were identified.

**Figure 6.**
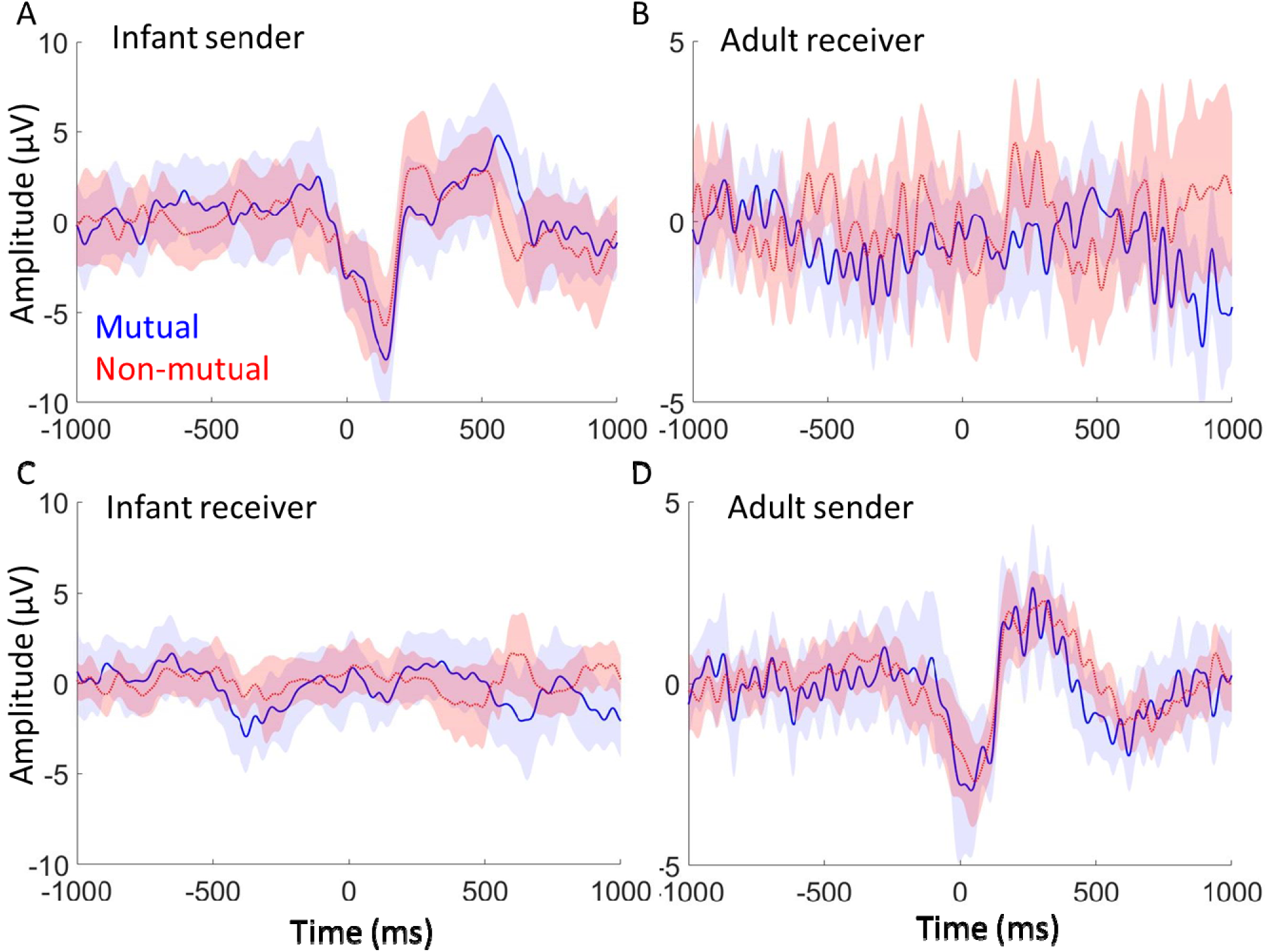
Event-related potentials time-locked to naturally occurring mutual and non-mutual gaze onsets. A) Infant ERP relative to onsets of infant sender mutual gaze and non-mutual gaze onsets. B) Adult ERP relative to adult receiver mutual and non-mutual gaze onsets. C) Infant ERP relative to onsets of infant receiver mutual and non-mutual gaze onsets. D) Adult ERP relative to adult sender mutual and non-mutual gaze onsets. For each the shaded area indicates 95% confidence intervals; thicker lines indicate grand average waveforms. Additional topoplots can be found in SM 7.

We then compared ERP amplitudes between mutual and non-mutual gaze onsets. As the ERP amplitudes were non-significant over baseline, relative to receiver mutual and non-mutual gaze onsets we focused our comparison on sender mutual vs non-mutual gaze onsets. The results of the paired samples t-test indicated no statistically significant differences after correcting for multiple comparisons. This was consistent for all three components; P1 (p=0.66 for the infant data and p=0.04 for the adult data; uncorrected p-values), N170/N290 (p=0.61 for infants and p=0.45 for adults) and P300/P400 (p=0.21 for infants, p=0.59 in adults).

Throughout the event locked analyses careful attention was paid to what activity reflected genuine neural responses and what was related to artifact. To investigate this in more detail we performed additional analyses (see SM section 2) in which we compared the senders’ neural responses pre and post artifact cleaning and compared activity over frontal vs occipital electrodes. The results of this analysis suggested that, despite the timing of the peak of the ITC occurring before the onset of gaze, it is unlikely that these findings are driven by the eye movement artifact itself, but rather the resulting neural response. In order to further test the sensitivity of our paradigm, we also compared sender neural responses between looks to object vs looks to partner gaze (see SM section 1). The results of this analysis suggested that our paradigm differentiates neural responses to face vs object looks, consistent with the results from previous ERP studies.

### 4.4. Intra-brain event locked analysis – ITC

Lastly, we examined the possibility that the onset of mutual gaze could act as synchronisation triggers to concomitantly reset the phase of the sender and receiver’s ongoing neural oscillations.

Figures 7 shows the results of the event-locked ITC analysis, comparing sender and receiver mutual and non-mutual gaze onsets for infant and adults separately. We first tested whether ITC values, over occipital electrodes, and frequencies 2-18 Hz, significantly exceeded baseline values generated from a permutation procedure (full details can be found in SM section 4). The permutation analysis indicated that ITC in the pre-gaze onset window was significantly higher than baseline for sender mutual (Figure 4 A and D) and non-mutual (Figure 4 E and H) gaze onsets, in both parents and infants, but not for receiver mutual (Figure 4 B and C) and non-mutual (Figure 4 F and G) gaze onsets in either parents or infants.

**Figure 7.**
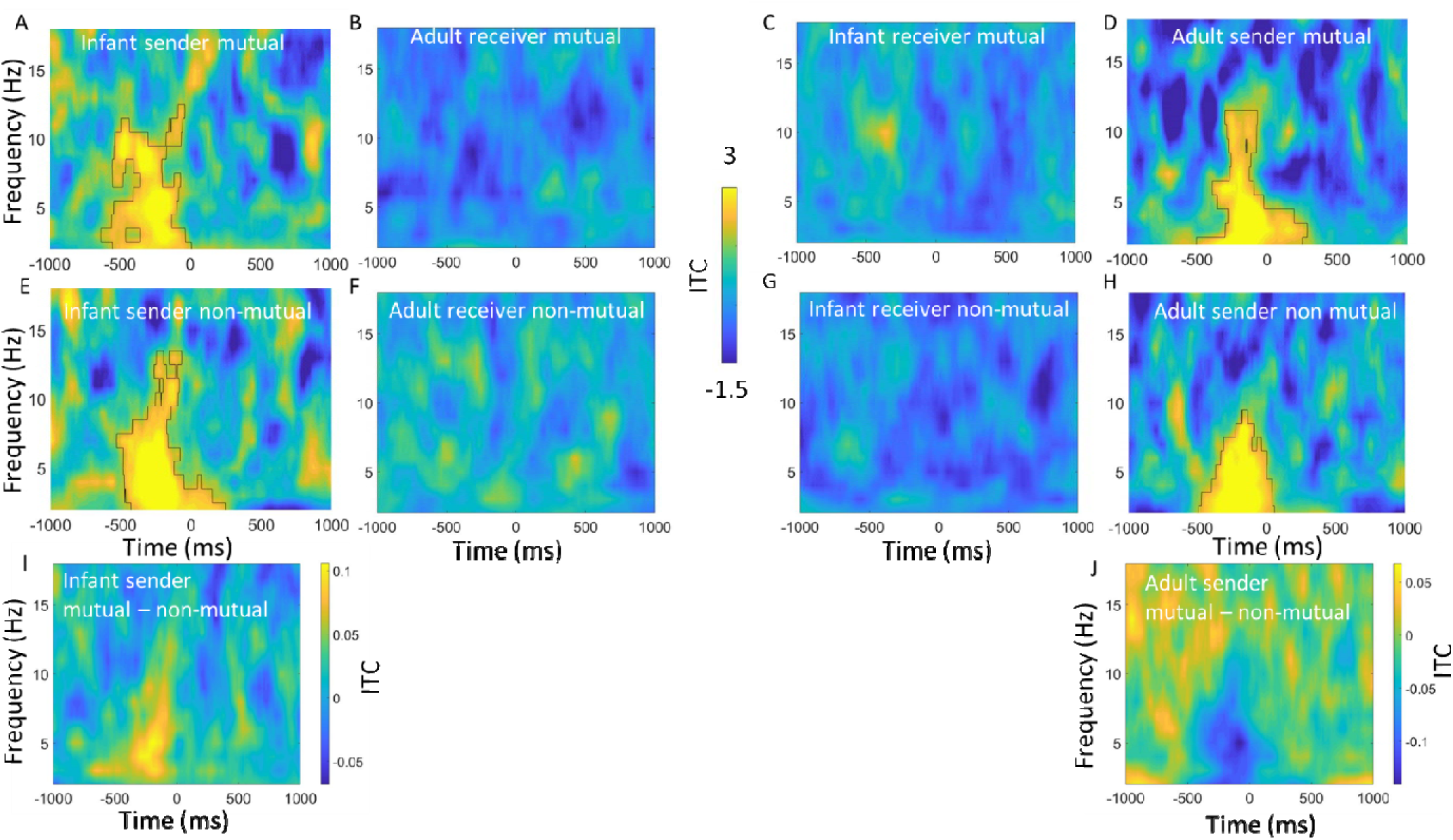
Z-scored inter-trial phase coherence time-locked to naturally occurring mutual and non-mutual gaze onsets. A) Infant ITC relative to infant sender mutual gaze onsets. B) Adult ITC relative to adult receiver mutual gaze onsets. C) Infant ITC relative to infant receiver mutual gaze onsets. D) Adult ITC relative to adult sender mutual gaze onsets. E) Infant ITC relative to infant sender non-mutual gaze onsets. F) Adult ITC relative to adult receiver non-mutual gaze onsets. G) Infant ITC relative to infant receiver non-mutual gaze onsets. H) Adult ITC relative to adult sender non-mutual gaze onsets. For all, black borders highlight activity that was significantly greater than baseline after cluster correction for multiple comparisons using p = .05. I) Difference plot between infant sender mutual vs non-mutual gaze onsets. J) Difference plot between adult sender mutual vs non-mutual gaze onsets. Hotter colours indicate more ITC for mutual vs non-mutual.

We then wanted to test whether ITC values were greater for mutual vs non-mutual gaze onsets. The results of the cluster-based permutation analysis indicated no statistically significant differences between looks to mutual vs non-mutual gaze in the sender’s brain activity in either parents or infants. The permutation analysis did reveal that ITC in the post gaze onset window was significantly greater relative to infant, but not adult receiver mutual vs non-mutual gaze onsets (peaking in Theta between 0-500ms post gaze onset, see S6).

## 5. Discussion

We took dual EEG recordings from parents and infants whilst they engaged in naturalistic free-flowing social interactions. Our data were analysed using cleaning and analysis procedures specially designed for naturalistic dual EEG data (Haresign et al., 2021; 2022). Since our analyses suggested that eye movement artifact cannot be completely removed from the EEG, we primarily compared sections of the data that were both identically time-locked to saccades, and therefore contain (presumably) an identical amount of eye movement artifact. The saccades were differentiated by the consequences of the saccade (either the saccade leads to mutual gaze, or not). Furthermore, we also compared activity only in the later parts of the ERP waveform after the first 100ms, when we were confident that no residual artifact remained. We considered three main sets of research questions:

### Mutual gaze and inter-brain synchrony

Our first set of research questions explored whether inter-brain synchrony was greater during mutual vs non-mutual gaze. Our results indicated that inter-brain synchrony did not significantly exceed baseline values for either mutual or non-mutual gaze. Further, comparing mutual vs non-mutual gaze, our results indicated that inter-brain synchrony was not greater during mutual vs non-mutual gaze, contrary to what we had hypothesised. These null findings were consistent across both frequencies and both measures of synchrony that we looked at (PLV and PDC), and across both our non-event and event locked analyses.

These results are inconsistent with previous studies that observed greater inter-brain synchrony during continuous (i.e., not relative to specific behaviours/ events within the interaction, but rather looking across all moments of a given behaviour during social interaction) moments of mutual vs non-mutual gaze. For example, in our previous paper we found increased inter-brain synchrony using PDC, in Theta and Alpha, over C3 and C4 electrodes in *N* = 29 *8*-month-olds (Leong et al., 2017). In the present study, we measured PDC and PLV across the same frequencies and electrodes in *N* = 55 *12*-month-olds.

Although we followed the same analytical techniques as Leong et al., 2017, we used different pre-processing techniques and a different (less structured, more naturalistic) paradigm, which could explain why our results differ. Firstly, the previous study featured an unfamiliar live adult singing nursery rhymes to an infant. Our present study, in contrast, featured primary caregivers interacting freely with the infant, using a puppet that they held in their hand. Infants’ sensitivity to novel interaction partners is well documented (Bushnell et al., 1989; de Haan & Nelson, 1997; 1999; Barry-Anwar et al., 2016; Hoehl et al., 2012). Therefore, one explanation for the positive effects of gaze type on inter-brain synchrony in our previous study could be due to the saliency of mutual gaze in the presence of an unfamiliar adult. Further research could investigate whether there are differences in inter-brain synchrony during infant-caregiver compared to infant-stranger interactions. Second, in our previous study, adults continuously sung nursery rhymes to the infants during the interactions, whereas in our present study they talked normally. As sung nursery rhymes are highly periodic (Suppanen et al., 2019) and evidence suggests that infant’s neural activity entrains to the temporal structure of these songs (Leong et al., 2017a; Attaheri et al., 2022), it could be that the regularity of the nursery rhymes introduced an external periodic stimulus into the environment that was driving the inter-brain entrainment (e.g., Perez et al., 2017). Here, mutual gaze might only enhance or maintain synchrony that is already established, by facilitating shared attention and therefore upregulating attention-enhanced neural synchrony. It will be important for future research to examine inter-brain synchrony in a variety of settings, ranging from very unstructured settings such as those used in the present study to more structured settings in which there are environmental stimuli with more regular and predictable inputs. Overall, these inconsistencies highlight the likely context-specific and localised nature of inter-brain synchrony, and further emphasise the importance of replication and standard data quality measures (Luck, 2021) when studying inter-brain dynamics (Holroyd, 2022).

### Phase resetting around gaze onsets

For our second set of research questions, we explored event-locked intra and inter-brain neural responses associated with mutual gaze onsets. Through this, we aimed to test our previously published hypothesis that concomitant phase resetting in the sender and the receiver’s brain at the onset of gaze may drive inter-brain synchrony (Leong et al., 2017; Wass et al., 2020). Overall, the results of our event-locked analyses are inconsistent with this idea. Contrary to our hypothesis, inter-brain synchrony did not significantly exceed baseline values for sender/ receiver mutual or non-mutual gaze onsets and was not significantly different between sender or receiver mutual vs non-mutual gaze onsets. Further, whilst we found that sender but not receiver mutual and non-mutual gaze onsets led to significant increases in ITC and amplitude (ERPs) over baseline, we did not find significant differences between sender or receiver mutual vs non-mutual gaze onsets. We did, however, find evidence for increases in ITC relative to sender mutual and non-mutual gaze onsets (section 3.4); but it is difficult to conclude that this represents phase resetting of brain oscillations. It could also be that changes in event locked amplitude/ power create the artifactual appearance of phase synchrony (Muthukumaraswamy et al., 2011) – a fact that the close correspondences we observed between ITC and event-locked changes in amplitude/power (see SM 2) would appear to support.

One possible driver of the sender neural responses could be residual eye movement artifact in our data. I<u>n</u> the supplementary analysis (SM section 3) we compare time-frequency power over frontal and occipital electrodes before and after ICA cleaning and report that ICA cleaning removed most, but not all, of the assumed artifactual activity associated with the eye movement- a conclusion consistent with our previous research (Haresign et al., 2021). This analysis also allowed us to identify that these artifacts are transient (∼100ms) and therefore only impacted the initial part of the ERP waveform. After the initial ∼+150-200ms we observed ERP components that look very similar to ERPs observed in traditional screen-based tasks (see Figure S1), with clear P1, N290 and P400 components. For added safety, however, our main analyses were based on comparing sections of the data that are both identically time-locked to saccades, and therefore contain an identical amount of eye movement artifact.

Overall, then, our results challenge the theory that phase resetting around key communicative signals such as mutual gaze is a mechanism through which inter-brain synchrony is achieved. Assuming that inter-brain synchrony according to more recent frameworks (Holroyd, 2022) is associated with mutual gaze. This points to the potential importance of other potential drivers of inter-brain synchrony, that future work should investigate in more detail – such as correlated changes in amplitude/ power or changes in oscillatory frequency independent of phase resetting (see Haresign et al., 2022 for a detailed discussion), and other more periodic behaviours (e.g., speech; Leong et al., 2017a; Attaheri et al., 2022).

### Re-examining the importance of ‘receiver’ mutual gaze in infant-caregiver social interaction

Our third aim was to test the hypothesis that infants are highly sensitive to ostensive signals during free-flowing social interactions with their caregivers. A number of influential papers (Farroni et al., 2002; 2004 Grossman et al., 2007; Senju & Johnson 2009; Csibra & Gergely, 2009) argue that, from shortly after birth, infants’ brains are sensitive to receiving ostensive signals, and that ‘sender’ communicative signals play a key role during naturalistic learning exchanges. However, as we noted these findings have not replicated well in developmental research; for example, Elsabbagh and colleagues (2009) or in research with adults (e.g., Watanbe et al., 2001; Taylor et al., 2001b; Watanbe et al., 2002; Itier et al., 2007; Conty et al., 2007; Ponkanen et al., 2011).

Contrary to expectations we found robust neural responses only in the senders’ (i.e., the agent initiating the gaze episode) and not in the receivers’ neural responses. This was true both for receiver non-mutual gaze onsets (where receivers were not looking at their partners and thus may have failed to detect their partners’ gaze shift), but also for the receiver-mutual condition (where receivers were directly gazing towards their partner at the time of the gaze shift). Evidence from adult ERP studies in which dynamic changes in gaze are simulated, through the presentation of a series of static images on a screen suggest that human adults are sensitive to ‘dynamic’ changes in gaze (Latinus et al., 2015; Stephani et al., 2020). However, these simulated changes in gaze are still far from the continuous way that gaze is processed during real social interactions, and it is likely that the effects observed in these studies are largely driven by more low-level properties of the simulation (e.g., retinal stimulation evoked by the presentation of a series of static images) rather than reflecting the actual processing of the gaze shift. When scrambled control images are presented in this same way this produces similar neural responses to those associated with processing simulated changes in gaze (Rossi et al., 2014). This suggests that whilst these studies do capture some neural mechanisms that are sensitive to moment-to-moment changes in the visual input from our environment, these studies do a poor job of simulating the continuous flow of gaze information that happens during real life social interaction. However, these studies do show some subtle neural sensitivity to changes in gaze orientation that perhaps we were unable to capture with the level of sensitivity afforded in our current approach. This raises basic questions over where, when and under what circumstances changes in a partner’s gaze during free-flowing social interactions impacts the neural activity of the receiver (the person viewing the gaze shift).

Again, one possible explanation for the inconsistencies between previous screen-based tasks and the present study is simply it is just a result of increased artifact through the use of a naturalistic paradigm. However, we note that: i) our ERPs show a close visual correspondence with ERPs observed in traditional ERP paradigms (see Figure 6 and Figure S1 also); ii) the overall measures of EEG data quality we reported show good quality data (see SM section 8); iii) we did replicate the findings from screen-based ERP research that infants show enhanced ERPs to images of faces vs objects (Guy et al., 2016; 2018; Peykarjou & Hoehl, 2013; Xie & Richards, 2016) (SM section 1); We observed statistically greater occipital ERP amplitudes for faces vs objects for the N290 component, but not for P1 or P400 components. Overall, then, we found changes in brain activity only in the person that initiated the episode of mutual gaze (the sender), and no changes in the recipient of the mutual gaze. This conclusion suggests a different account of the dyadic mechanisms involved in the processing of mutual gaze. In contrast to evidence suggesting that mutual gaze involves both infants and adults reciprocally influencing each other’s neural activity towards shared rhythms (Leong et al., 2017), we found changes in brain activity only in the person that initiated the episode of mutual gaze (the sender). This suggests that in this context, mutual gaze processing is more supported through more basic changes at the intra-brain level, which do not specifically affect dyadic neural mechanisms. For example, evidence suggests that eye movements lead to low frequency phase reorganisation in brain structures such as the hippocampus that are deeper than those that can be measured using scalp EEG (e.g., Hoffman et al., 2013). Eye movements may create transient increases in neural sensitivity within certain structures within an individual’s brain that support them in processing the new visual information (e.g., mutual gaze) (Klimesch et al., 2007).

## 6. Conclusion

We investigated the possibility that concomitant phase resetting in response to mutual gaze onsets during naturalistic infant-caregiver interactions might be a mechanism through which inter-brain synchrony is established. We found no evidence for changes in inter-brain synchrony around gaze onsets and no evidence to support our previously published suggestion that phase resetting in the sender and the receiver’s brain around mutual gaze onsets may be a mechanism through which inter-brain synchrony arises (Leong et al., 2017; Wass et al., 2020). Further, contrary to our prediction, we found that mutual gaze onsets associated with neural responses in ‘senders’, but not in ‘receivers’ brains. Overall, our study challenges current views on the importance of mutual gaze. It highlights the fact that we need pluralistic approaches to better understand early social cognition. And it highlights the importance of studying how infants perceive communicative signals during naturalistic interactions, and across different real-world contexts.

## Acknowledgements

We wish to thank Victoria Leong for contributing to task design, for advice and providing processing scripts for data analysis, and for reading multiple drafts of the manuscript. We also wish to thank members of the UEL BabyDev Lab for useful advice, and our experimental participants. This research was funded by Project Grant RPG-2018-281 from the Leverhulme Trust, by an ERC Marie Curie Fellowship JDIL 845859 and by ERC grant number ONACSA 853251.

## References

Attaheri, A., Choisdealbha, Á. N., Di Liberto, G. M., Rocha, S., Brusini, P., Mead, N., … & Goswami, U. (2022). Delta-and theta-band cortical tracking and phase-amplitude coupling to sung speech by infants. NeuroImage, 247, 118698.

Azzalini, D., Rebollo, I., & Tallon-Baudry, C. (2019). Visceral signals shape brain dynamics and cognition. Trends in cognitive sciences, 23(6), 488–509.

Baccalá, L. A., & Sameshima, K. (2001). Partial directed coherence: a new concept in neural structure determination. Biological cybernetics, 84(6), 463–474.

Barnett, L., & Seth, A. K. (2014). The MVGC multivariate Granger causality toolbox: a new approach to Granger-causal inference. Journal of neuroscience methods, 223, 50–68.

Barrett, A. B., Murphy, M., Bruno, M. A., Noirhomme, Q., Boly, M., Laureys, S., & Seth, A. K. (2012). Granger causality analysis of steady-state electroencephalographic signals during propofol-induced anaesthesia. PloS one, 7(1), e29072.

Barry-Anwar, R. A., Burris, J. L., Estes, K. G., & Rivera, S. M. (2017). Caregivers and strangers: The influence of familiarity on gaze following and learning. Infant Behavior and Development, 46, 46–58.

Bart Krekelberg (2022). bayesFactor (https://github.com/klabhub/bayesFactor), GitHub. Retrieved February 28, 2022.

Begus, K., Gliga, T., & Southgate, V. (2014). Infants learn what they want to learn: Responding to infant pointing leads to superior learning. PloS one, 9(10), e108817.

Begus, K., Gliga, T., & Southgate, V. (2016). Infants’ preferences for native speakers are associated with an expectation of information. Proceedings of the National Academy of Sciences, 113(44), 12397–12402.

Billings, R. J. (1989). The origin of the occipital lambda wave in man. Electroencephalography and clinical neurophysiology, 72(2), 95–113.

Bloom, L., Lightbown, P., Hood, L., Bowerman, M., Maratsos, M., & Maratsos, M. P. (1975). Structure and variation in child language. Monographs of the society for Research in Child Development, 1–97.

Biau, E., Torralba, M., Fuentemilla, L., de Diego Balaguer, R., & Soto-Faraco, S. (2015). Speaker’s hand gestures modulate speech perception through phase resetting of ongoing neural oscillations. Cortex, 68, 76–85.

Burgess, A. P. (2013). On the interpretation of synchronization in EEG hyperscanning studies: a cautionary note. Frontiers in human neuroscience, 7, 881.

Busch, N. A., Dubois, J., & VanRullen, R. (2009). The phase of ongoing EEG oscillations predicts visual perception. Journal of neuroscience, 29(24), 7869–7876.

Bushnell, I. W. R., Sai, F., & Mullin, J. T. (1989). Neonatal recognition of the mother’s face. British journal of developmental psychology, 7(1), 3–15.

Calderone, D. J., Lakatos, P., Butler, P. D., & Castellanos, F. X. (2014). Entrainment of neural oscillations as a modifiable substrate of attention. Trends in cognitive sciences, 18(6), 300–309.

Carpenter, M., Nagell, K., Tomasello, M., Butterworth, G., & Moore, C. (1998). Social cognition, joint attention, and communicative competence from 9 to 15 months of age. Monographs of the society for research in child development, i-174.

Çetinçelik, M., Rowland, C. F., & Snijders, T. M. (2021). Do the eyes have it? A systematic review on the role of eye gaze in infant language development. Frontiers in psychology, 11, 3627.

Cohen, M. X. (2014). Analyzing neural time series data: theory and practice. MIT press.

Conte, S., Richards, J. E., Guy, M. W., Xie, W., & Roberts, J. E. (2020). Face-sensitive brain responses in the first year of life. NeuroImage, 211, 116602.

Conty, L., N’Diaye, K., Tijus, C., & George, N. (2007). When eye creates the contact! ERP evidence for early dissociation between direct and averted gaze motion processing. Neuropsychologia, 45(13), 3024–3037.

Csibra, G. (2010). Recognizing communicative intentions in infancy. Mind & Language, 25(2), 141–168.

Csibra, G., & Gergely, G. (2009). Natural pedagogy. Trends in cognitive sciences, 13(4), 148–153.

Delorme, A., & Makeig, S. (2004). EEGLAB: an open source toolbox for analysis of single-trial EEG dynamics including independent component analysis. Journal of neuroscience methods, 134(1), 9–21.

De Haan, M., & Nelson, C. A. (1997). Recognition of the mother’s face by sixLmonthLold infants: A neurobehavioral study. Child development, 68(2), 187–210.

De Haan, M., & Nelson, C. A. (1999). Brain activity differentiates face and object processing in 6-month-old infants. Developmental psychology, 35(4), 1113.

De Jaegher, H. (2009). Social understanding through direct perception? Yes, by interacting. Consciousness and cognition, 18(2), 535–542.

De Munck, J. C., & Bijma, F. (2010). How are evoked responses generated? The need for a unified mathematical framework. Clinical neurophysiology: official journal of the International Federation of Clinical Neurophysiology, 121(2), 127–129.

Dimigen, O. (2020). Optimizing the ICA-based removal of ocular EEG artifacts from free viewing experiments. NeuroImage, 207, 116117.

Dimigen, O. (2021). Fixation-related potentials in total darkness. Journal of Vision, 21(9), 2931–2931.

Dimigen, O., Valsecchi, M., Sommer, W., & Kliegl, R. (2009). Human microsaccade-related visual brain responses. Journal of Neuroscience, 29(39), 12321–12331.

Dimigen, O., Sommer, W., Hohlfeld, A., Jacobs, A. M., & Kliegl, R. (2011). Coregistration of eye movements and EEG in natural reading: analyses and review. Journal of experimental psychology: General, 140(4), 552.

Doelling, K. B., & Assaneo, M. F. (2021). Neural oscillations are a start toward understanding brain activity rather than the end. PLoS Biology, 19(5), e3001234.

Donnellan, E., Bannard, C., McGillion, M. L., Slocombe, K. E., & Matthews, D. (2020). Infants’ intentionally communicative vocalizations elicit responses from caregivers and are the best predictors of the transition to language: A longitudinal investigation of infants’ vocalizations, gestures and word production. Developmental Science, 23(1), e12843.

Elsabbagh, M., Volein, A., Csibra, G., Holmboe, K., Garwood, H., Tucker, L., … & Johnson, M. H. (2009). Neural correlates of eye gaze processing in the infant broader autism phenotype. Biological psychiatry, 65(1), 31–38.

Faes, L., Erla, S., Porta, A., & Nollo, G. (2013). A framework for assessing frequency domain causality in physiological time series with instantaneous effects. Philosophical Transactions of the Royal Society A: Mathematical, Physical and Engineering Sciences, 371(1997), 20110618.

Farroni, T., Csibra, G., Simion, F., & Johnson, M. H. (2002). Eye contact detection in humans from birth. Proceedings of the National academy of sciences, 99(14), 9602–9605.

Farroni, T., Massaccesi, S., Pividori, D., & Johnson, M. H. (2004). Gaze following in newborns. Infancy, 5(1), 39–60.

Faul, F., Erdfelder, E., Lang, A. G., & Buchner, A. (2007). G* Power 3: A flexible statistical power analysis program for the social, behavioral, and biomedical sciences. Behavior research methods, 39(2), 175–191.

Feldman, R., Magori-Cohen, R., Galili, G., Singer, M., & Louzoun, Y. (2011). Mother and infant coordinate heart rhythms through episodes of interaction synchrony. Infant Behavior and Development, 34(4), 569–577.

Ferguson, B., & Lew-Williams, C. (2016). Communicative signals support abstract rule learning by 7-month-old infants. Scientific reports, 6(1), 1–7.

Franchak, J. M., Kretch, K. S., Soska, K. C., & Adolph, K. E. (2011). HeadLmounted eye tracking: A new method to describe infant looking. Child development, 82(6), 1738–1750.

Gaarder, K., Krauskopf, J., Graf, V., Kropfl, W., & Armington, J. C. (1964). Averaged brain activity following saccadic eye movement. Science, 146(3650), 1481–1483.

Georgieva, S., Lester, S., Noreika, V., Yilmaz, M. N., Wass, S., & Leong, V. (2020). Toward the understanding of topographical and spectral signatures of infant movement artifacts in naturalistic EEG. Frontiers in neuroscience, 14, 352.

Giraud, A. L., & Poeppel, D. (2012). Cortical oscillations and speech processing: emerging computational principles and operations. Nature neuroscience, 15(4), 511–517.

Grossmann, T., & Johnson, M. H. (2007). The development of the social brain in human infancy. European Journal of Neuroscience, 25(4), 909–919.

Guy, M. W., Zieber, N., & Richards, J. E. (2016). The cortical development of specialized face processing in infancy. Child Development, 87(5), 1581–1600.

Guy, M. W., Richards, J. E., Tonnsen, B. L., & Roberts, J. E. (2018). Neural correlates of face processing in etiologically-distinct 12-month-old infants at high-risk of autism spectrum disorder. Developmental Cognitive Neuroscience, 29, 61–71.

Hains, S. M., & Muir, D. W. (1996). Infant sensitivity to adult eye direction. Child development, 67(5), 1940–1951.

Hamilton, A. F. D. C. (2021). Hyperscanning: beyond the hype. Neuron, 109(3), 404–407.

Haresign, I. M., Phillips, E., Whitehorn, M., Noreika, V., Jones, E. J. H., Leong, V., & Wass, S. V. (2021). Automatic classification of ICA components from infant EEG using MARA. Developmental cognitive neuroscience, 52, 101024.

Haresign, I. M., Phillips, E. A. M., Whitehorn, M., Goupil, L., Noreika, V., Leong, V., & Wass, S. V. (2022). Measuring the temporal dynamics of inter-personal neural entrainment in continuous child-adult EEG hyperscanning data. Developmental cognitive neuroscience, 54, 101093.

Henry, M. J., Herrmann, B., & Obleser, J. (2014). Entrained neural oscillations in multiple frequency bands comodulate behavior. Proceedings of the National Academy of Sciences, 111(41), 14935–14940.

Hirsch, J., Zhang, X., Noah, J. A., & Ono, Y. (2017). Frontal temporal and parietal systems synchronize within and across brains during live eye-to-eye contact. Neuroimage, 157, 314–330.

Hoehl, S., & Markova, G. (2018). Moving developmental social neuroscience toward a second-person approach. PLoS Biology, 16(12), e3000055.

Hoehl, S., Wahl, S., Michel, C., & Striano, T. (2012). Effects of eye gaze cues provided by the caregiver compared to a stranger on infants’ object processing. Developmental Cognitive Neuroscience, 2(1), 81–89.

Hoehl, S., Wiese, L., & Striano, T. (2008). Young infants’ neural processing of objects is affected by eye gaze direction and emotional expression. PLoS One, 3(6), e2389.

Hoffman, K. L., Dragan, M. C., Leonard, T. K., Micheli, C., Montefusco-Siegmund, R., & Valiante, T. A. (2013). Saccades during visual exploration align hippocampal 3–8 Hz rhythms in human and non-human primates. Frontiers in systems neuroscience, 7, 43.

Holroyd, C. B. (2022). Interbrain synchrony: on wavy ground. Trends in Neurosciences.

Itier, R. J., Alain, C., Sedore, K., & McIntosh, A. R. (2007). Early face processing specificity: it’s in the eyes!. Journal of cognitive neuroscience, 19(11), 1815–1826.

Jensen, O., Gips, B., Bergmann, T. O., & Bonnefond, M. (2014). Temporal coding organized by coupled Alpha and gamma oscillations prioritize visual processing. Trends in neurosciences, 37(7), 357–369.

Jones, E. J., Goodwin, A., Orekhova, E., Charman, T., Dawson, G., Webb, S. J., & Johnson, M. H. (2020). Infant EEG theta modulation predicts childhood intelligence. Scientific reports, 10(1), 1–10.

Kayhan, E., Matthes, D., Haresign, I. M., Bánki, A., Michel, C., Langeloh, M., … & Hoehl, S. (2022). DEEP: A dual EEG pipeline for developmental hyperscanning studies. Developmental cognitive neuroscience, 54, 101104.

Kazai, K., & Yagi, A. (2003). Comparison between the lambda response of eye-fixation-related potentials and the P100 component of pattern-reversal visual evoked potentials. Cognitive, Affective, & Behavioral Neuroscience, 3(1), 46–56.

Konvalinka, I., & Roepstorff, A. (2012). The two-brain approach: how can mutually interacting brains teach us something about social interaction?. Frontiers in human neuroscience, 6, 215.

Krakauer, J. W., Ghazanfar, A. A., Gomez-Marin, A., MacIver, M. A., & Poeppel, D. (2017). Neuroscience needs behavior: correcting a reductionist bias. Neuron, 93(3), 480–490.

Klimesch, W., Sauseng, P., & Hanslmayr, S. (2007). EEG Alpha oscillations: the inhibition– timing hypothesis. Brain research reviews, 53(1), 63–88.

Lachaux, J. P., Rodriguez, E., Martinerie, J., & Varela, F. J. (1999). Measuring phase synchrony in brain signals. Human brain mapping, 8(4), 194–208.

Latinus, M., Love, S. A., Rossi, A., Parada, F. J., Huang, L., Conty, L., … & Puce, A. (2015). Social decisions affect neural activity to perceived dynamic gaze. Social cognitive and affective neuroscience, 10(11), 1557–1567.

Leong, V., Byrne, E., Clackson, K., Georgieva, S., Lam, S., & Wass, S. (2017). Speaker gaze increases information coupling between infant and adult brains. Proceedings of the National Academy of Sciences, 114(50), 13290–13295.

Lester, B. M., Hoffman, J., & Brazelton, T. B. (1985). The rhythmic structure of mother-infant interaction in term and preterm infants. Child development, 15–27.

Luft, C., Zioga, I., Giannopoulos, A., Di Bona, G., Civilini, A., Latora, V., & Mareschal, I. (2021). Social synchronisation of brain activity by eye-contact.

Luck, S. J., Stewart, A. X., Simmons, A. M., & Rhemtulla, M. (2021). Standardized measurement error: A universal metric of data quality for averaged eventLrelated potentials. Psychophysiology, 58(6), e13793.

Marshall, P. J., Young, T., & Meltzoff, A. N. (2011). Neural correlates of action observation and execution in 14LmonthLold infants: An eventLrelated EEG desynchronization study. Developmental science, 14(3), 474–480.

Mathewson, K. E., Gratton, G., Fabiani, M., Beck, D. M., & Ro, T. (2009). To see or not to see: prestimulus α phase predicts visual awareness. Journal of Neuroscience, 29(9), 2725–2732.

Mathewson, K. E., Fabiani, M., Gratton, G., Beck, D. M., & Lleras, A. (2010). Rescuing stimuli from invisibility: Inducing a momentary release from visual masking with pre-target entrainment. Cognition, 115(1), 186–191.

Mathewson, K. E., Lleras, A., Beck, D. M., Fabiani, M., Ro, T., & Gratton, G. (2011). Pulsed out of awareness: EEG Alpha oscillations represent a pulsed-inhibition of ongoing cortical processing. Frontiers in psychology, 2, 99.

Mathewson, K. E., Prudhomme, C., Fabiani, M., Beck, D. M., Lleras, A., & Gratton, G. (2012). Making waves in the stream of consciousness: entraining oscillations in EEG Alpha and fluctuations in visual awareness with rhythmic visual stimulation. Journal of cognitive neuroscience, 24(12), 2321–2333.

McHugh, M. L. (2012). Interrater reliability: the kappa statistic. Biochemia medica, 22(3), 276–282.

Muthukumaraswamy, S. D., & Singh, K. D. (2011). A cautionary note on the interpretation of phase-locking estimates with concurrent changes in power. Clinical neurophysiology: official journal of the International Federation of Clinical Neurophysiology, 122(11), 2324–2325.

Novembre, G., & Iannetti, G. D. (2021). Hyperscanning alone cannot prove causality. Multibrain stimulation can. Trends in Cognitive Sciences, 25(2), 96–99.

Pan, Y., Novembre, G., & Olsson, A. (2021). The interpersonal neuroscience of social learning. Perspectives on Psychological Science, 17456916211008429.

Peykarjou, S., & Hoehl, S. (2013). Three-month-olds’ brain responses to upright and inverted faces and cars. Developmental Neuropsychology, 38(4), 272–280.

Pérez, A., Carreiras, M., & Duñabeitia, J. A. (2017). Brain-to-brain entrainment: EEG interbrain synchronization while speaking and listening. Scientific reports, 7(1), 1–12.

Piazza, E. A., Hasenfratz, L., Hasson, U., & Lew-Williams, C. (2020). Infant and adult brains are coupled to the dynamics of natural communication. Psychological Science, 31(1), 6–17.

Phillips, E., Goupil, L., Haresign, I. M., Bruce-Gardyne, E., Csolsim, F. A., Whitehorn, M., … & Wass, S. (2021). Proactive or reactive? Neural oscillatory insight into the leader-follower dynamics of early infant-caregiver interaction.

Plöchl, M., Ossandón, J. P., & König, P. (2012). Combining EEG and eye tracking: identification, characterization, and correction of eye movement artifacts in electroencephalographic data. Frontiers in human neuroscience, 6, 278.

Pönkänen, L. M., Alhoniemi, A., Leppänen, J. M., & Hietanen, J. K. (2011). Does it make a difference if I have an eye contact with you or with your picture? An ERP study. Social cognitive and affective neuroscience, 6(4), 486–494.

Rayson, H., Bonaiuto, J. J., Ferrari, P. F., Chakrabarti, B., & Murray, L. (2019). Building blocks of joint attention: Early sensitivity to having one’s own gaze followed. Developmental cognitive neuroscience, 37, 100631.

Redcay, E., & Schilbach, L. (2019). Using second-person neuroscience to elucidate the mechanisms of social interaction. Nature Reviews Neuroscience, 20(8), 495–505.

Rossi, A., Parada, F. J., Kolchinsky, A., & Puce, A. (2014). Neural correlates of apparent motion perception of impoverished facial stimuli: a comparison of ERP and ERSP activity. NeuroImage, 98, 442–459.

Ruzzoli, M., Torralba, M., Fernández, L. M., & Soto-Faraco, S. (2019). The relevance of Alpha phase in human perception. Cortex, 120, 249–268.

Schilbach, L. (2010). A second-person approach to other minds. Nature Reviews Neuroscience, 11(6), 449–449.

Schilbach, L., Timmermans, B., Reddy, V., Costall, A., Bente, G., Schlicht, T., & Vogeley, K. (2013). Toward a second-person neuroscience 1. Behavioral and brain sciences, 36(4), 393–414.

Schroeder, C. E., & Lakatos, P. (2009). Low-frequency neuronal oscillations as instruments of sensory selection. Trends in neurosciences, 32(1), 9–18.

Southgate, V., Chevallier, C., & Csibra, G. (2010). SeventeenLmonthLolds appeal to false beliefs to interpret others’ referential communication. Developmental science, 13(6), 907–912.

Stephani, T., Driller, K. K., Dimigen, O., & Sommer, W. (2020). Eye contact in active and passive viewing: Event-related brain potential evidence from a combined eye tracking and EEG study. Neuropsychologia, 143, 107478.

Suppanen, E., Huotilainen, M., & Ylinen, S. (2019). Rhythmic structure facilitates learning from auditory input in newborn infants. Infant Behavior and Development, 57, 101346.

Szymanski, C., Pesquita, A., Brennan, A. A., Perdikis, D., Enns, J. T., Brick, T. R., … & Lindenberger, U. (2017). Teams on the same wavelength perform better: Inter-brain phase synchronization constitutes a neural substrate for social facilitation. Neuroimage, 152, 425–436.

Symons, L. A., Hains, S. M., & Muir, D. W. (1998). Look at me: Five-month-old infants’ sensitivity to very small deviations in eye-gaze during social interactions. Infant Behavior and Development, 21(3), 531–536.

Taylor, M. J., Edmonds, G. E., McCarthy, G., & Allison, T. (2001). Eyes first! Eye processing develops before face processing in children. Neuroreport, 12(8), 1671–1676.

Thickbroom, G. W., Knezevic, W., Carroll, W. M., & Mastaglia, F. L. (1991). Saccade onset and offset lambda waves: relation to pattern movement visually evoked potentials. Brain research, 551(1-2), 150–156.

Thut, G., Miniussi, C., & Gross, J. (2012). The functional importance of rhythmic activity in the brain. Current Biology, 22(16), R658–R663.

van der Velde, B., White, T., & Kemner, C. (2021). The emergence of a theta social brain network during infancy. NeuroImage, 240, 118298.

Wass, S., & Goupil, L. (2022). Studying the developing brain in real-world contexts: moving from castles in the air to castles on the ground.

Wass, S. V., Whitehorn, M., Haresign, I. M., Phillips, E., & Leong, V. (2020). Interpersonal neural entrainment during early social interaction. Trends in cognitive sciences, 24(4), 329–342.

Wass, S. V., Noreika, V., Georgieva, S., Clackson, K., Brightman, L., Nutbrown, R., … & Leong, V. (2018). Parental neural responsivity to infants’ visual attention: How mature brains influence immature brains during social interaction. PLoS biology, 16(12), e2006328.

Watanabe, S., Kakigi, R., & Puce, A. (2001). Occipitotemporal activity elicited by viewing eye movements: a magnetoencephalographic study. Neuroimage, 13(2), 351–363.

Watanabe, S., Miki, K., & Kakigi, R. (2002). Gaze direction affects face perception in humans. Neuroscience letters, 325(3), 163–166.

Werker, J. F., Pons, F., Dietrich, C., Kajikawa, S., Fais, L., & Amano, S. (2007). Infant-directed speech supports phonetic category learning in English and Japanese. Cognition, 103(1), 147–162.

Werker, J. F., & Yeung, H. H. (2005). Infant speech perception bootstraps word learning. Trends in cognitive sciences, 9(11), 519–527.

Wohltjen, S., & Wheatley, T. (2021). Eye contact marks the rise and fall of shared attention in conversation. Proceedings of the National Academy of Sciences, 118(37).

Van Diepen, R. M., Cohen, M. X., Denys, D., & Mazaheri, A. (2015). Attention and temporal expectations modulate power, not phase, of ongoing Alpha oscillations. Journal of cognitive neuroscience, 27(8), 1573–1586.

VanRullen, R. (2016). Perceptual cycles. Trends in cognitive sciences, 20(10), 723–735.

Vecera, S. P., & Johnson, M. H. (1995). Gaze detection and the cortical processing of faces: Evidence from infants and adults. Visual cognition, 2(1), 59–87.

Nguyen, T., Schleihauf, H., Kayhan, E., Matthes, D., Vrtička, P., & Hoehl, S. (2021). Neural synchrony in mother–child conversation: Exploring the role of conversation patterns. Social Cognitive and Affective Neuroscience, 16(1-2), 93–102.

Yu, C., & Smith, L. B. (2013). Joint attention without gaze following: Human infants and their parents coordinate visual attention to objects through eye-hand coordination. PloS one, 8(11), e79659.

Xie, W., Mallin, B. M., & Richards, J. E. (2018). Development of infant sustained attention and its relation to EEG oscillations: an EEG and cortical source analysis study. Developmental Science, 21(3), e12562.

Xie, W., & Richards, J. E. (2016). Effects of interstimulus intervals on behavioral, heart rate, and eventLrelated potential indices of infant engagement and sustained attention. Psychophysiology, 53(8), 1128–1142.

