## Supplementary materials for "Gaze onsets during naturalistic infant-caregiver interaction associate with ‘sender’ but not ‘receiver’ neural responses, and do not lead to changes in inter-brain synchrony"

### 1. *Intra brain analysis – ERPs to faces vs objects*

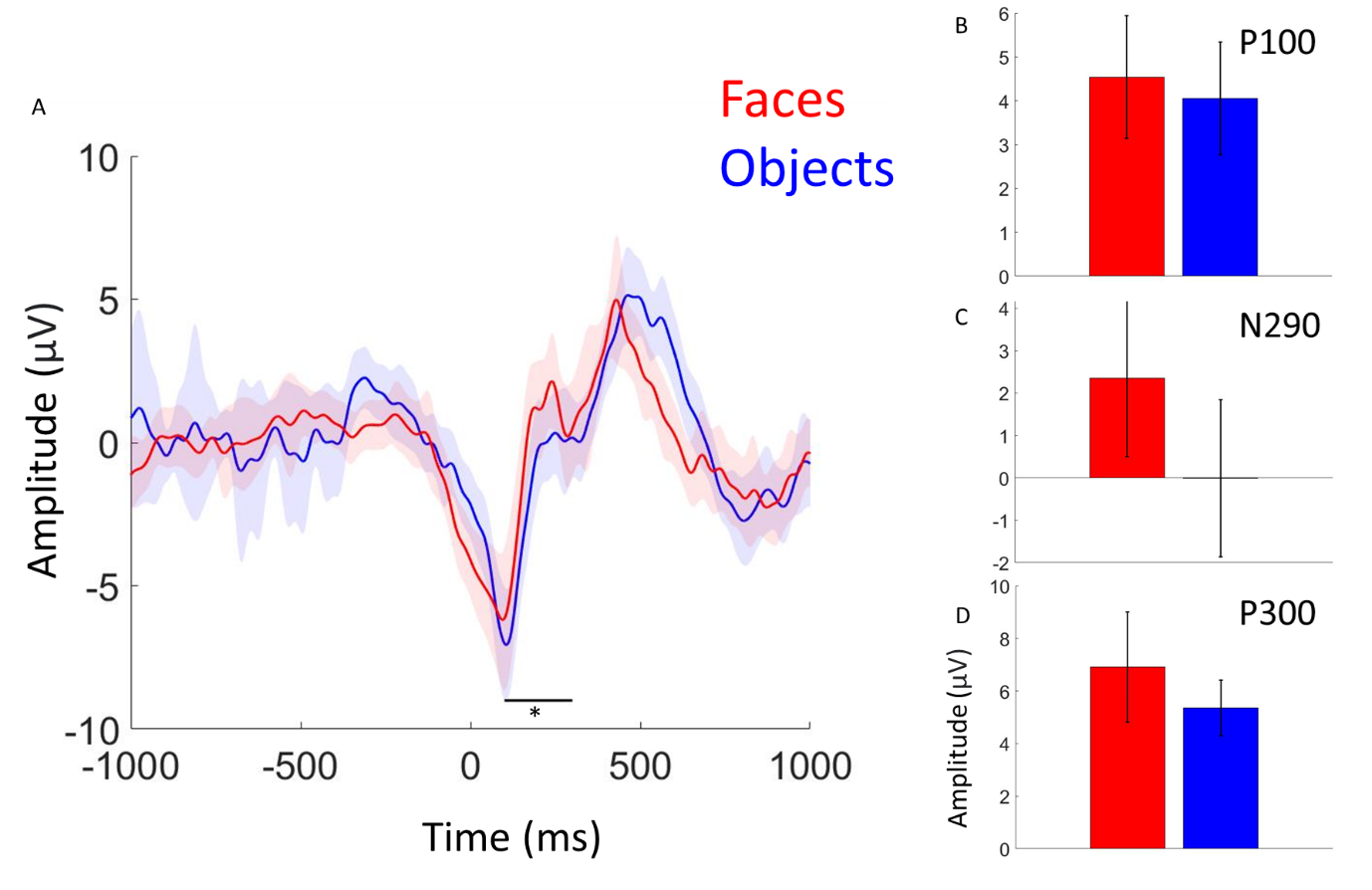

*Figure S1. Event-related potentials time-locked to naturally occurring partner (face) gaze and object gaze onsets. A) Infant occipital ERPs relative to onsets of infant looks to their partners face vs infant looks to object. Shaded areas indicate 95% confidence intervals, thicker lines indicate grand average waveforms and black line indicates time points at which ERP amplitudes for faces vs objects were significantly greater at p = .05.B) Grand average amplitudes for P100 component for faces vs objects. C) Grand average amplitudes for N290 component for faces vs objects. D) Grand average amplitudes for P300 component for faces vs objects. Bar charts are for visual purposes only to show the direction of the difference. Error bars indicate 95% confidence intervals.*

Figure S1 shows the results of the ERP analysis, comparing infant initiated partner and object looks. As an additional analysis to our comparison between mutual and non-mutual gaze we wanted to replicate previous findings (e.g., Conte et al., 2020) that show greater infant ERP amplitudes to faces vs objects. We observed statistically greater occipital ERP amplitudes for faces vs objects for the N290 component (*p =* 0.03), but not for P1 or P400 components, *p =* 0.14/ *p =* 0.50 respectively. Note these represent uncorrected p values. Our results suggested that our paradigm differentiates neural responses to face vs object looks, consistent with the results from previous ERP studies.

### 2. *Intra brain event locked analysis - Power*

It is known that changes in spectral power resulting from evoked neural responses can give the appearance of increased phase-locking/ resetting, due to changes in signal to noise ratios and error associated with estimating phase (Muthukumaraswamy et al., 2011). Power was obtained as the square of the absolute values derived from the wavelet convolution procedure described in section 3.4 of the main text. Time-frequency power was baseline normalised (decibel normalised) using activity in the -1000 to -700ms time window and averaged over trials.

Differences between mutual and non-mutual gaze onsets in spectral power were assessed following the same statistical procedure as detailed in section 2.15 of the main text, using a cluster-based correction for multiple comparisons with an alpha value of .05. We found no significant differences in spectral power between gaze types after correction for multiple comparisons (see Figure S2). However, it is important to note the visual similarities in the time-frequency characteristics between the observed event locked ITC (see Figure 4) and power changes around gaze onsets. The significance of this is discussed within section 5.2 of
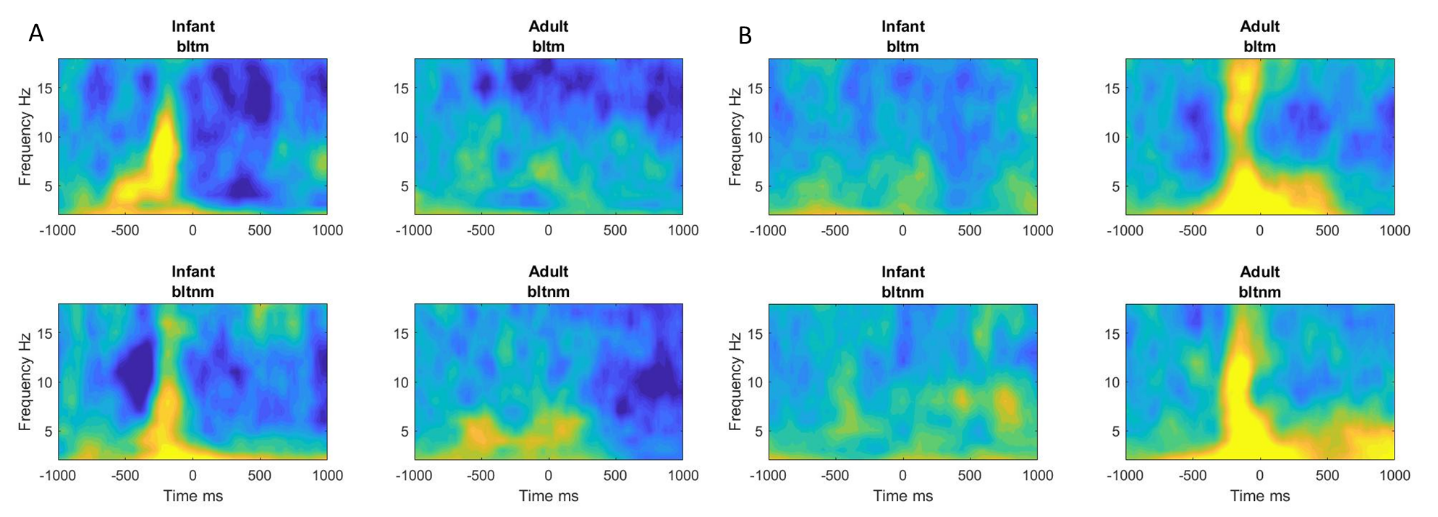
the main text.

*Figure S2. Occipital power time-locked to naturally occurring mutual and non-mutual gaze onsets. A) Infant and adult occipital power relative to infant sender/ adult receiver looks to mutual (bltm) and non-mutual gaze (bltnm) onsets. B) Infant and adult occipital power relative to adult sender/ infant looks to mutual (mltm) and non-mutual gaze (mltnm) onsets.*

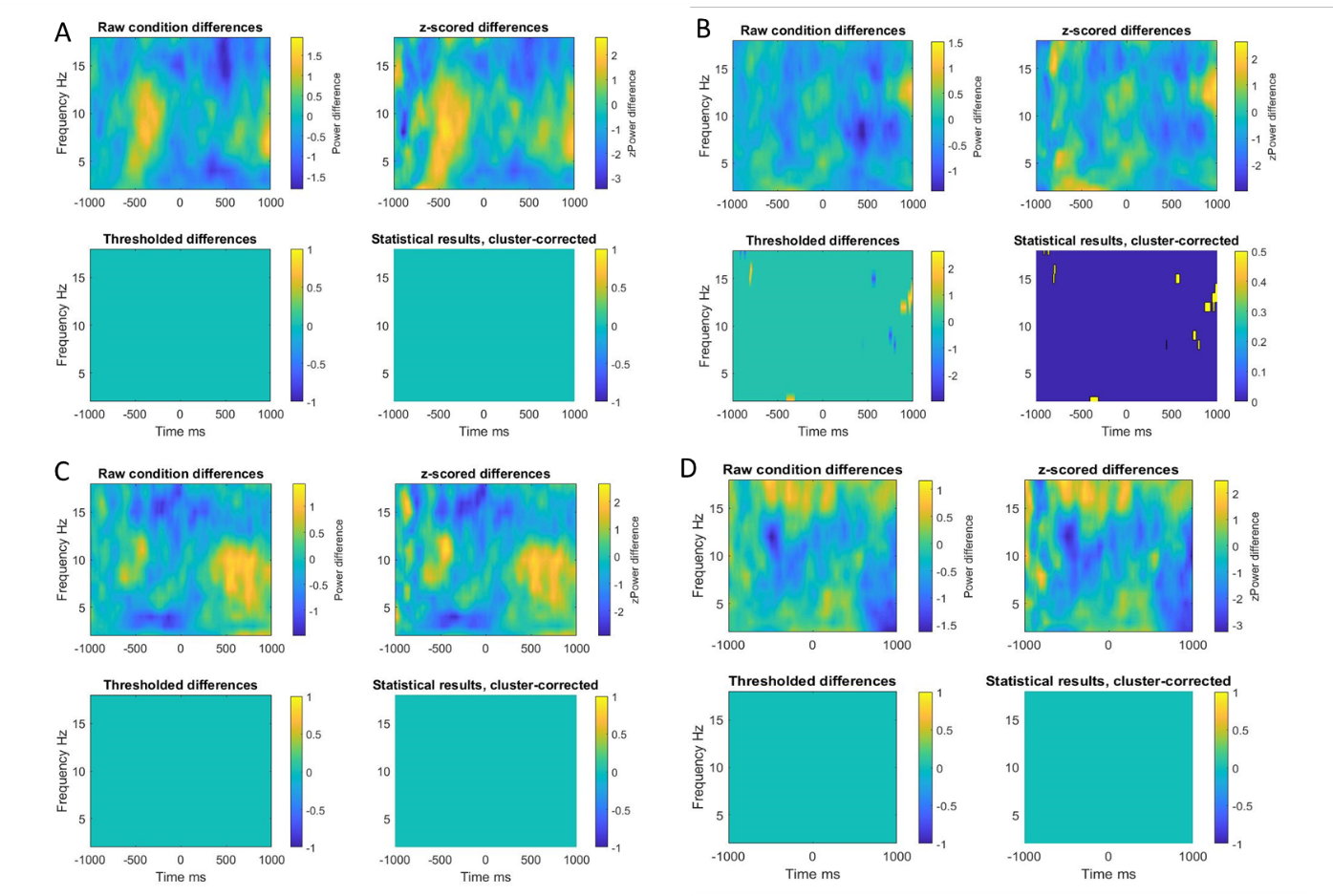

*Figure S3. Result of cluster-based permutation procedure for event locked power for gaze type A) Result of permutation for infant occipital power comparing infant sender/adult receiver looks to mutual vs non-mutual. B) Result of permutation for adult occipital power comparing infant sender/ adult receiver mutual vs non-mutual gaze onsets. C) Result of permutation for infant occipital power comparing adult sender/ infant receiver mutual vs non-mutual gaze onsets. D) Result of permutation for adult occipital power comparing adult sender/ infant receiver mutual vs non-mutual gaze onsets. For each, the top left plot is the raw differences where hotter (more yellow) colours indicate more power for mutual vs non-mutual. The top right plot shows z scored differences. The bottom left plot shows thresholded differences (at p = .05) before cluster correction for multiple comparisons. The bottom right shows the final results of cluster-based permutation statistics after correction for multiple comparisons.*

### 3. *Timing of ITC effect*

From Figure 4 of the main text, it can be seen that the peak of the ITC effect precedes the onset of the look. Here we conducted additional analysis to explore the possibility that the observed ITC effects could be a result of the eye movement artifacts and not the neural response associated with the processing of the new gaze information. To do this we first examined how the ITC effects varied topographically. Specifically, we looked at whether ITC was stronger over frontal (averaged over electrodes Fp1, Fp2, Af3, Af4) vs occipital (averaged over electrodes O1, O2, Oz, Po3, Po4) channels. From Figure S3 it can be seen that the ITC is weaker over frontal than occipital channels, suggesting that the occipital ITC is more strongly associated with the neural response rather than the eye movement artifact as we observed similar topographical patterns for amplitude and power (e.g., see SM section 2).

However, it is possible that our pre-processing procedure was bias toward removing frontal activity and therefore could have created the topographical differences we observed for ITC. In our data, we attempted to remove eye-movement artifacts using ICA decomposition, and an automated procedure was adopted for judging which components should be removed (Marriott Haresign et al., 2021). Though ICA algorithms are not biased towards removing specific types of artifacts, eye movement artifacts are often more stereotyped and easier to identify in an ICA decomposition (Chaumon et al., 2015), which could have led to biases in the systems judgement of which ICA components should be removed. Therefore, we also checked the topographical distribution of the observed ITC effects in data that had not been cleaned using ICA. Here we found consistent with our main findings, ITC was strongest over occipital electrodes. Overall these data suggest that, despite the timing of the peak of the ITC occurring before the onset of gaze, it is unlikely that these findings are driven by the eye movement artifact itself, but rather the resulting neural response.

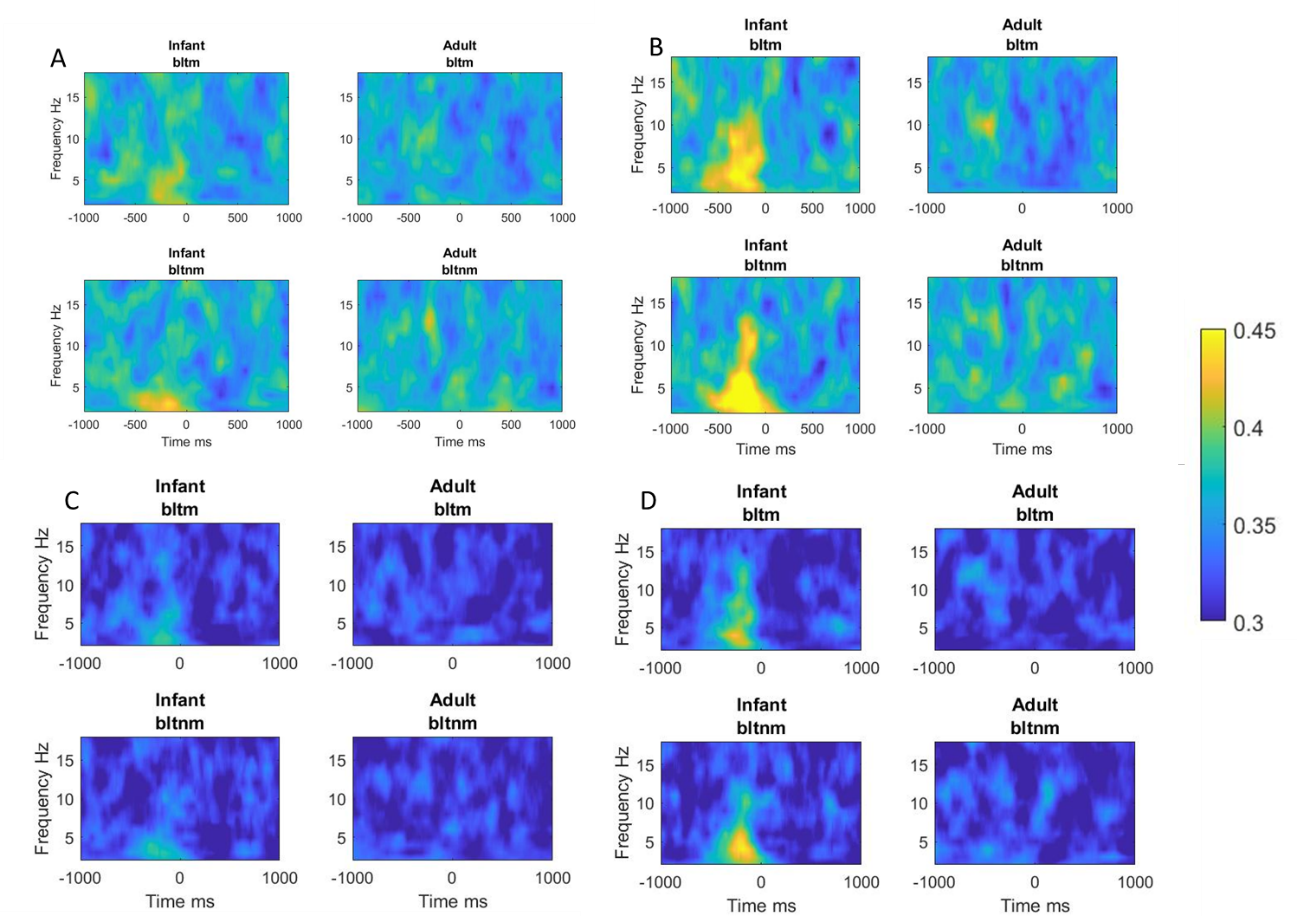

*Figure S4. Event-locked ITC time-locked to naturally occurring mutual and non-mutual gaze onsets before and after ICA cleaning. A) Infant and adult frontal ITC relative to infant sender/ adult receiver mutual and non-mutual gaze onsets post-ICA. B) Infant and adult occipital ITC relative to infant sender/ adult receiver mutual and non-mutual gaze onsets post-ICA. C) Infant and adult frontal ITC relative to adult sender/ infant receiver mutual and non-mutual gaze onsets pre-ICA. D) Infant and adult occipital ITC relative to adult sender/ infant receiver mutual and non-mutual gaze onsets pre-ICA.*

### 4. *Intra-brain event locked analysis ITC – Results of permutation procedure*

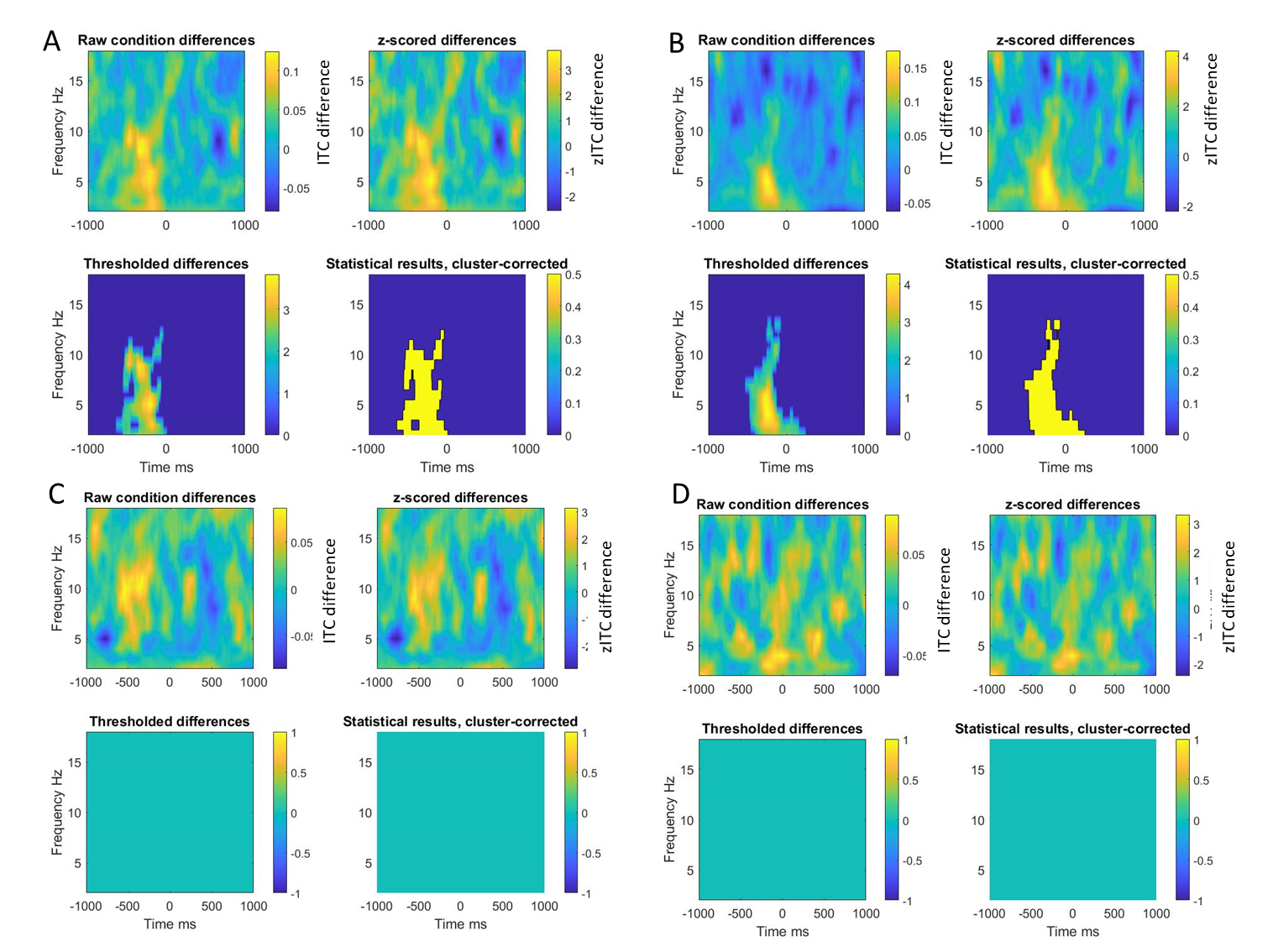

*Figure S5. Result of cluster-based permutation procedure for event locked ITC relative to baseline. A) Result of cluster-based permutation procedure for infant occipital ITC relative to infant sender/adult receiver mutual gaze onsets. B) Result of cluster-based permutation procedure for infant occipital ITC relative to infant sender/adult receiver non-mutual gaze onsets. C) Result of cluster-based permutation procedure for infant occipital ITC relative to adult sender/ infant receiver mutual gaze onsets. D) Result of cluster-based permutation procedure for infant occipital ITC relative to adult sender/infant receiver non-mutual gaze onsets. For each, the top left plot is the raw differences where hotter (more yellow) colours indicate more ITC for looks to mutual vs non-mutual. The top right plot shows z scored differences. The bottom left plot shows thresholded differences (at p=0.05) before cluster correction for multiple comparisons. The bottom right shows the final results of cluster-based permutation statistics after correction for multiple comparisons.*

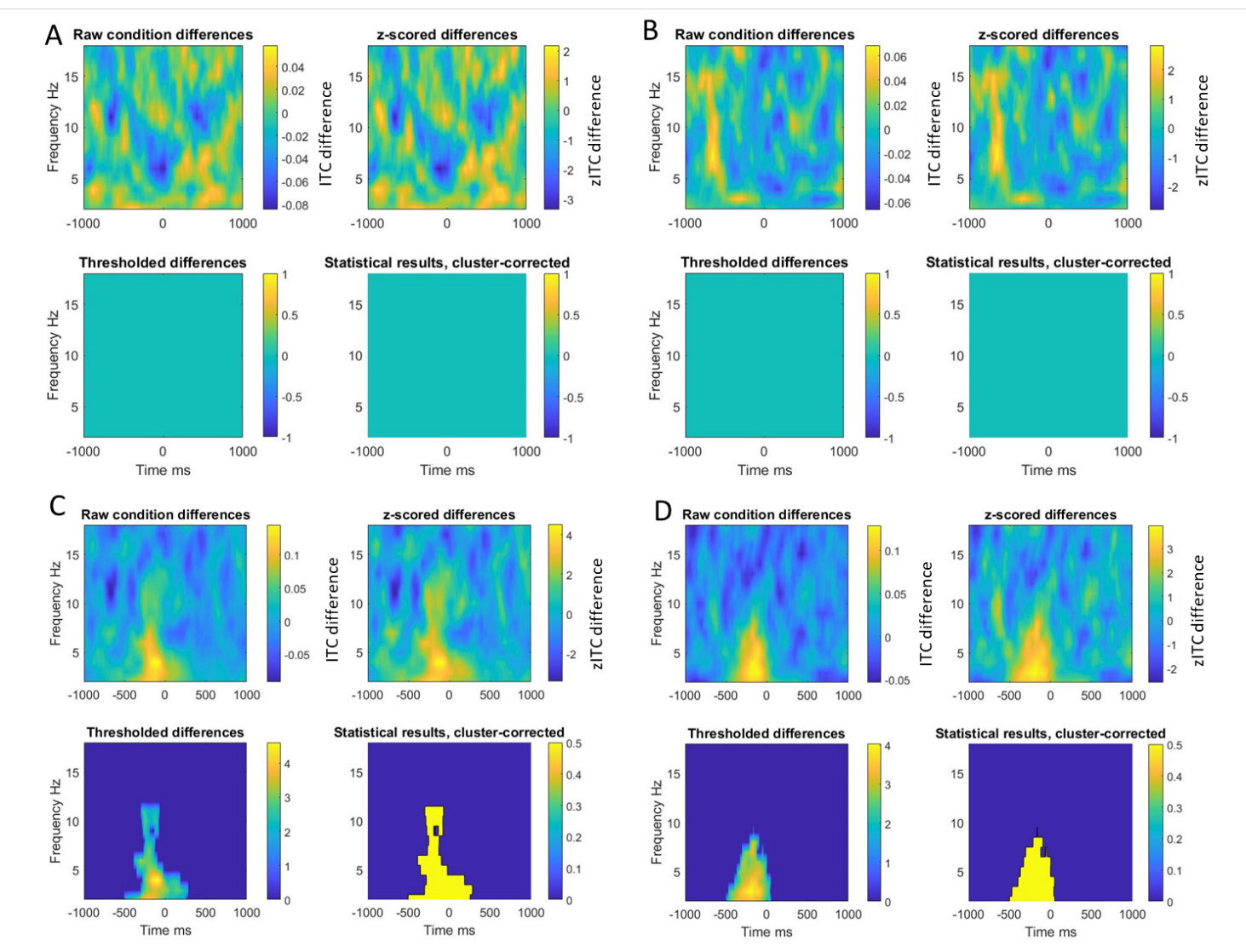

*Figure S6. Result of cluster-based permutation procedure for event locked ITC relative to baseline. A) Result of cluster-based permutation procedure for adult occipital ITC relative to infant sender/ adult receiver mutual gaze onsets. B) Result of cluster-based permutation procedure for adult occipital ITC relative to infant sender/ adult receiver non-mutual gaze onsets. C) Result of cluster-based permutation procedure for adult occipital ITC relative to adult sender/ infant receiver mutual gaze onsets. D) Result of cluster-based permutation procedure for adult occipital ITC relative to adult sender/ infant receiver non-mutual gaze onsets. For each, the top left plot is the raw differences where hotter (more yellow) colours indicate more ITC for looks to mutual vs non-mutual. The top right plot shows z scored differences. The bottom left plot shows thresholded differences (at p=0.05) before cluster correction for multiple comparisons. The bottom right shows the final results of cluster-based permutation statistics after correction for multiple comparisons.*

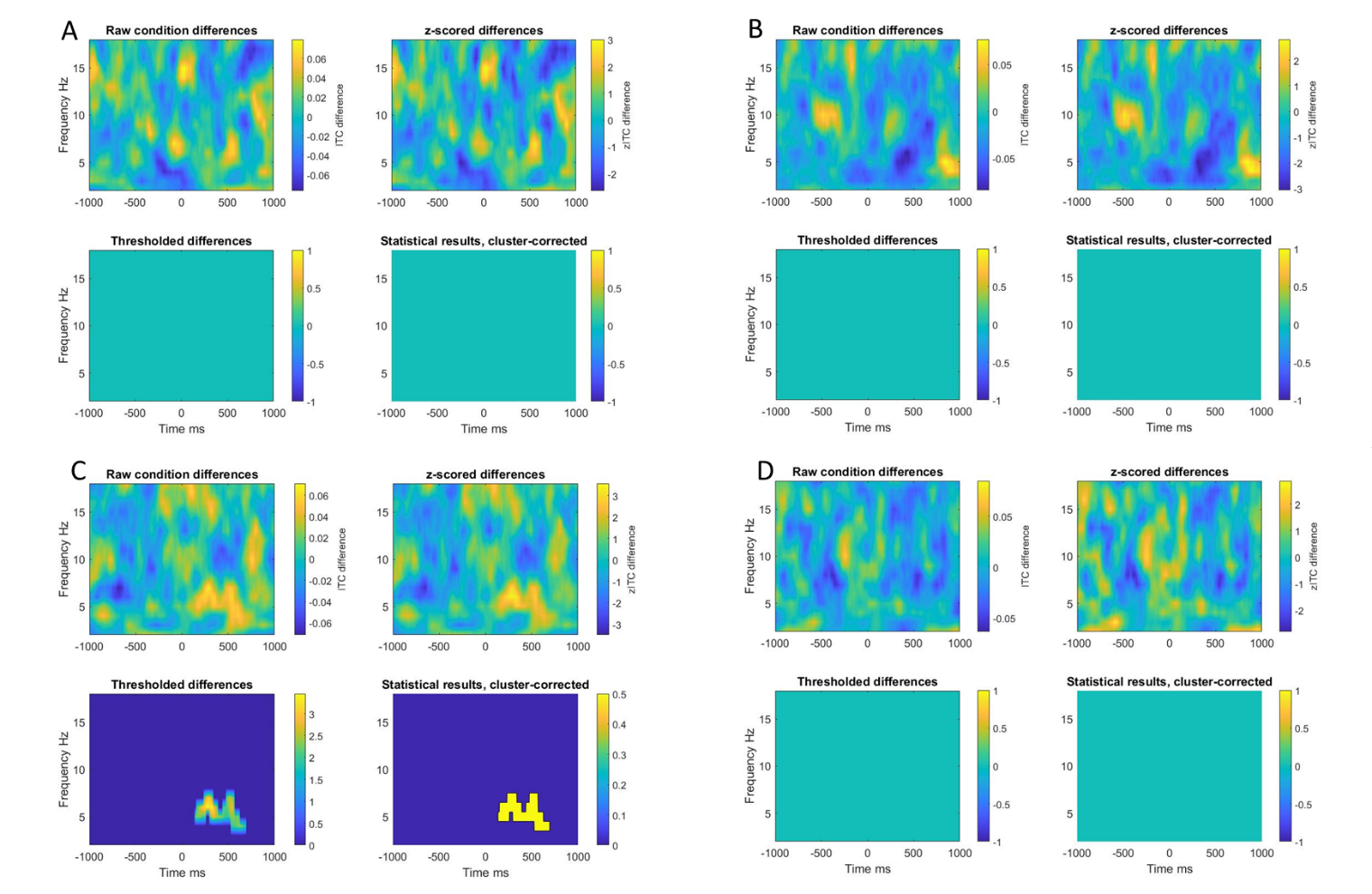

*Figure S7. Result of cluster-based permutation procedure for event locked ITC for effect of gaze type A) Result of cluster-based permutation procedure for infant occipital ITC comparing infant sender mutual vs non-mutual gaze onsets. B) Result of cluster-based permutation procedure for adult occipital ITC comparing infant sender mutual vs non-mutual gaze onset. C) Result of cluster-based permutation procedure for infant occipital ITC comparing adult sender mutual vs non-mutual gaze onsets. D) Result of cluster-based permutation procedure for adult occipital ITC comparing adult sender mutual vs non-mutual gaze onsets. For each, the top left plot is the raw differences where hotter (more yellow) colours indicate more ITC for looks to mutual vs non-mutual. The top right plot shows z scored differences. The bottom left plot shows thresholded differences (at p=0.05) before cluster correction for multiple comparisons. The bottom right shows the final results of cluster-based permutation statistics after correction for multiple comparisons.*

### 5. *Inter-brain event locked analysis PLV and PDC – Results of permutation procedure*

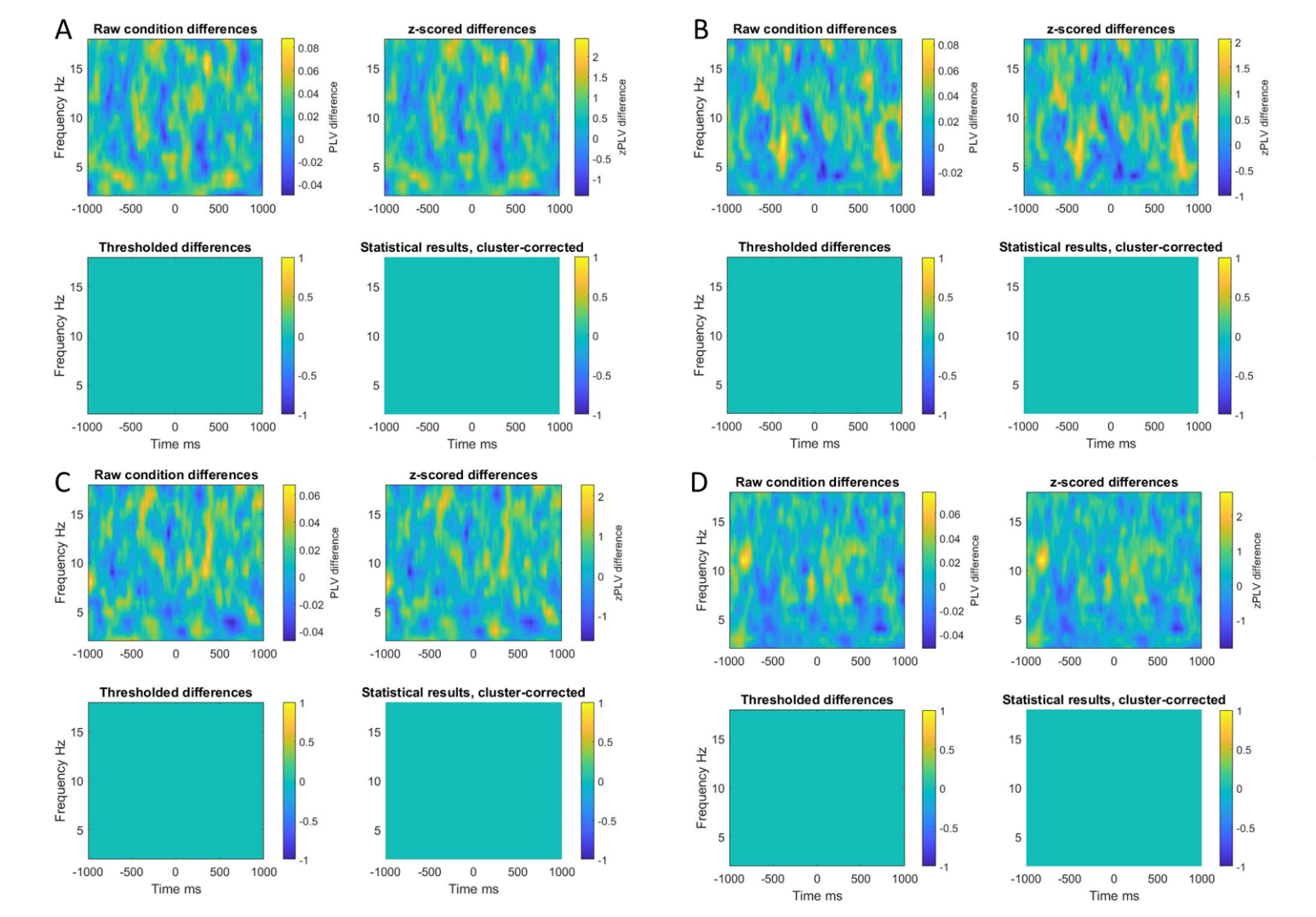

*Figure S8. Result of cluster-based permutation procedure for event locked PLV relative to baseline. A) Result of cluster-based permutation procedure for occipital PLV relative to infant sender mutual gaze onsets. B) Result of cluster-based permutation procedure for occipital PLV relative to infant sender non-mutual gaze onsets. C) Result of cluster-based permutation procedure for occipital PLV relative to adult sender mutual gaze onsets. D) Result of cluster-based permutation procedure for occipital PLV relative to adult sender non-mutual gaze onsets. For each, the top left plot is the raw differences where hotter (more yellow) colours indicate more PLV for looks to mutual vs non-mutual. The top right plot shows z scored differences. The bottom left plot shows thresholded differences (at p=0.05) before cluster correction for multiple comparisons. The bottom right shows the final results*
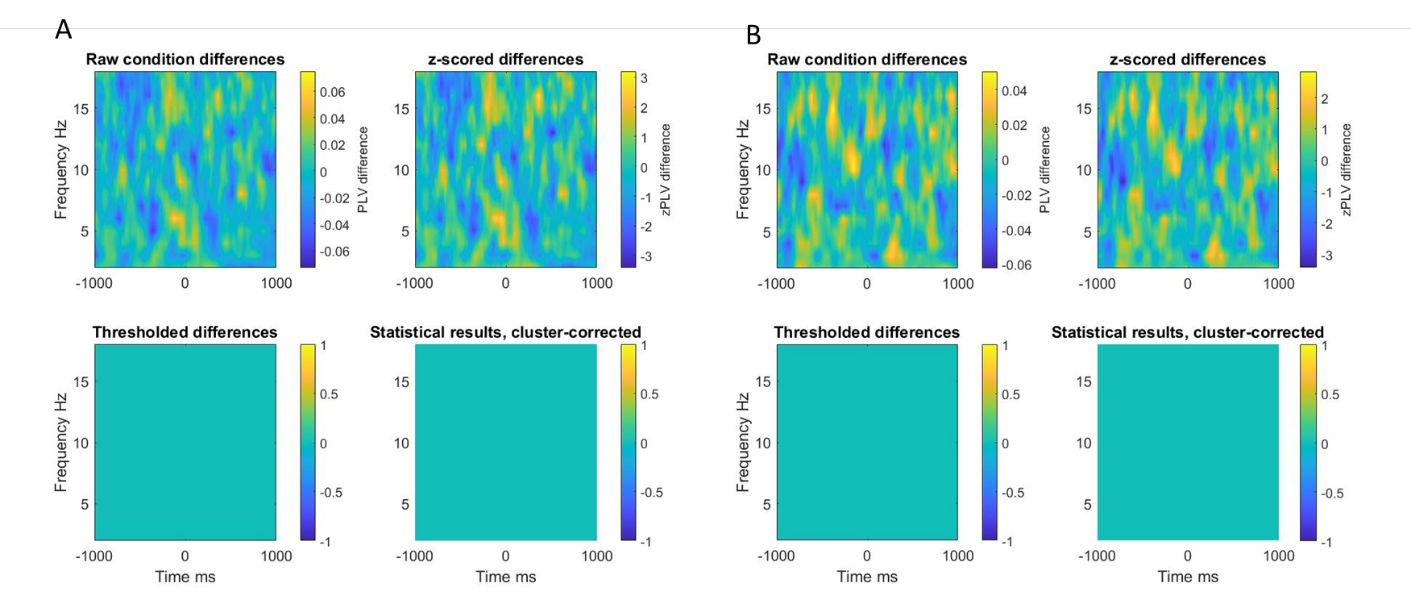
*of cluster-based permutation statistics after correction for multiple comparisons.*

*Figure S9. Result of cluster-based permutation procedure for PLV over occipital electrodes. A) shows difference between infant sender/ adult reciever looks to mutual vs non-mutual gaze onsets. B) shows difference between adult sender/ infant recevier mutual vs non-mutual gaze onsets (hotter, more yellow colours indicate more PLV for mutual vs non-mutual gaze).*

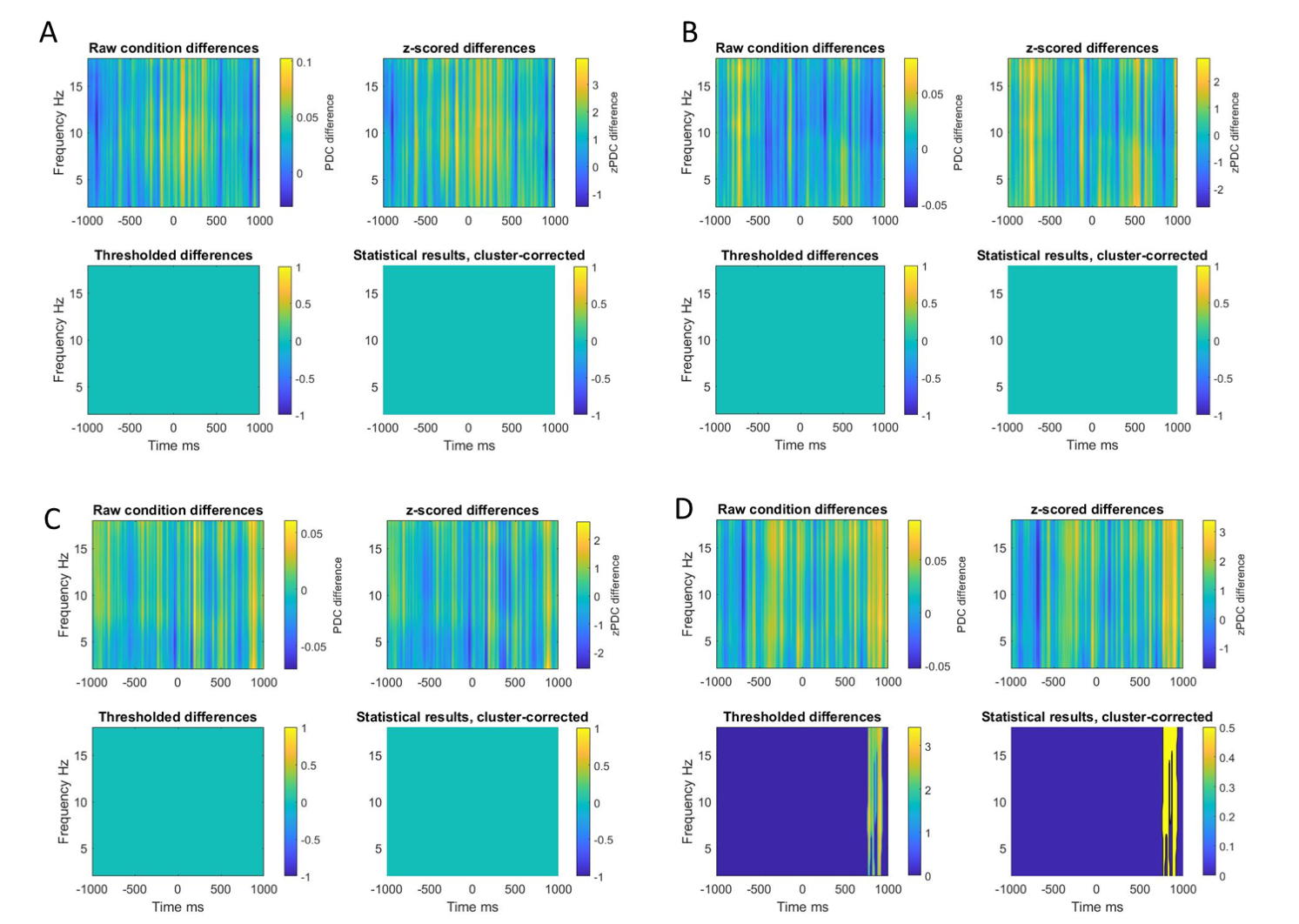

*Figure S10. Result of cluster-based permutation procedure for event locked PDC relative to baseline. A) Result of cluster-based permutation procedure for occipital I🡪A PDC relative to infant sender/ adult receiver mutual gaze onsets. B) Result of cluster-based permutation procedure for I🡪A occipital PLV relative to infant sender/ adult receiver non-mutual gaze onsets. C) Result of cluster-based permutation procedure for occipital A🡪I PDC relative to infant sender/ adult receiver mutual gaze onsets. D) Result of cluster-based permutation procedure for A🡪I occipital PLV relative to infant sender/ adult receiver non-mutual gaze onsets. For each, the top left plot is the raw differences where hotter (more yellow) colours indicate more PDC for mutual vs non-mutual. The top right plot shows z scored differences. The bottom left plot shows thresholded differences (at p=0.05) before cluster correction for multiple comparisons. The bottom right shows the final results of cluster-based permutation statistics after correction for multiple comparisons. I🡪A means infant to adult influence. A🡪I means adult to infant influence.*

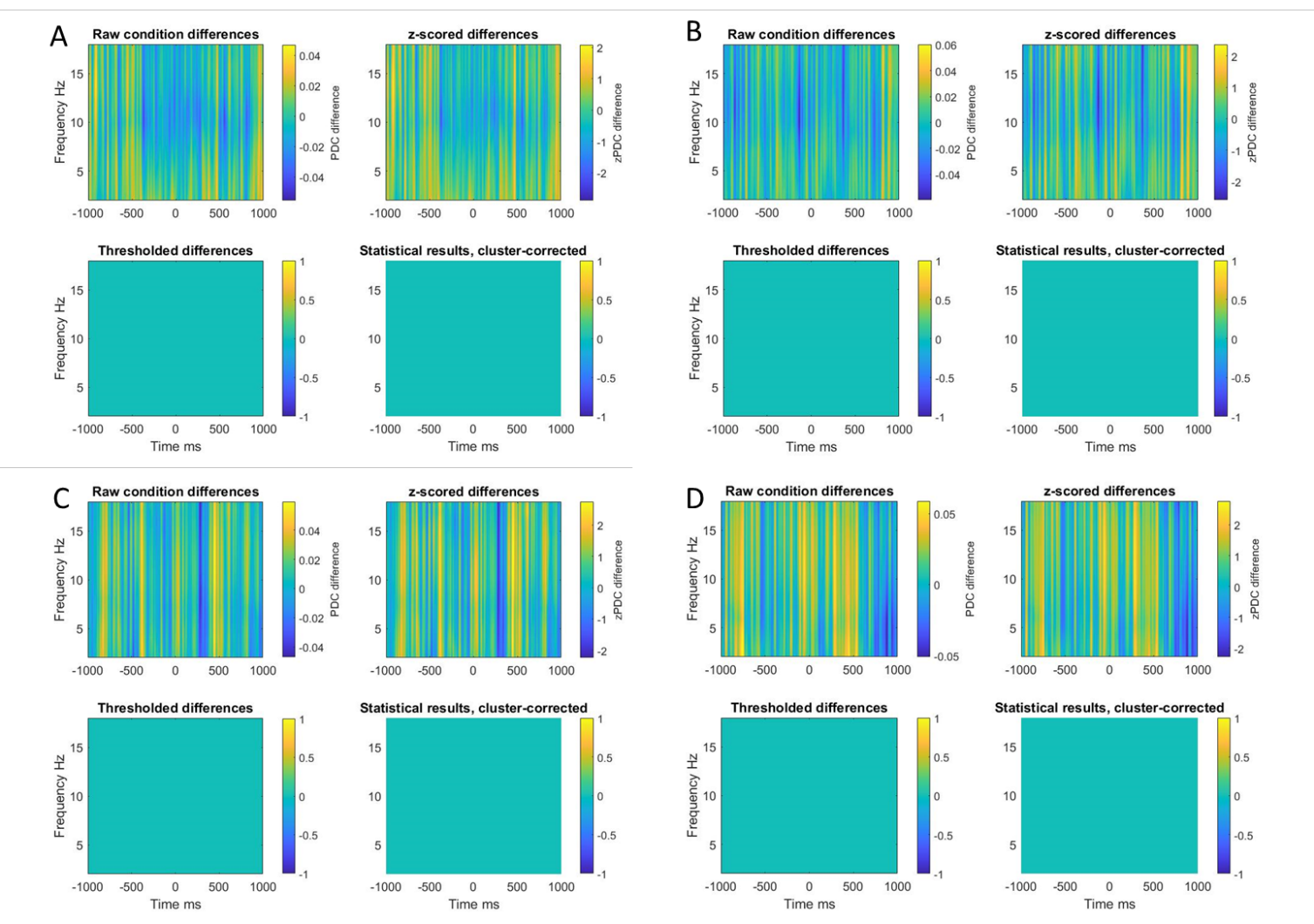

*Figure S11. Result of cluster-based permutation procedure for event locked PDC relative to baseline. A) Result of cluster-based permutation procedure for occipital I🡪A PDC relative to adult sender/ infant receiver mutual gaze onsets. B) Result of cluster-based permutation procedure for I🡪A occipital PLV relative to adult sender/ infant receiver non-mutual gaze onsets. C) Result of cluster-based permutation procedure for occipital A🡪I PDC relative to adult sender/ infant receiver mutual gaze onsets. D) Result of cluster-based permutation procedure for A🡪I occipital PLV relative to adult sender/ infant receiver non-mutual gaze onsets. For each, the top left plot is the raw differences where hotter (more yellow) colours indicate more PDC for mutual vs non-mutual. The top right plot shows z scored differences. The bottom left plot shows thresholded differences (at p=0.05) before cluster correction for multiple comparisons. The bottom right shows the final results of cluster-based permutation statistics after correction for multiple comparisons. I🡪A means infant to adult influence. A🡪I means adult to infant influence.*

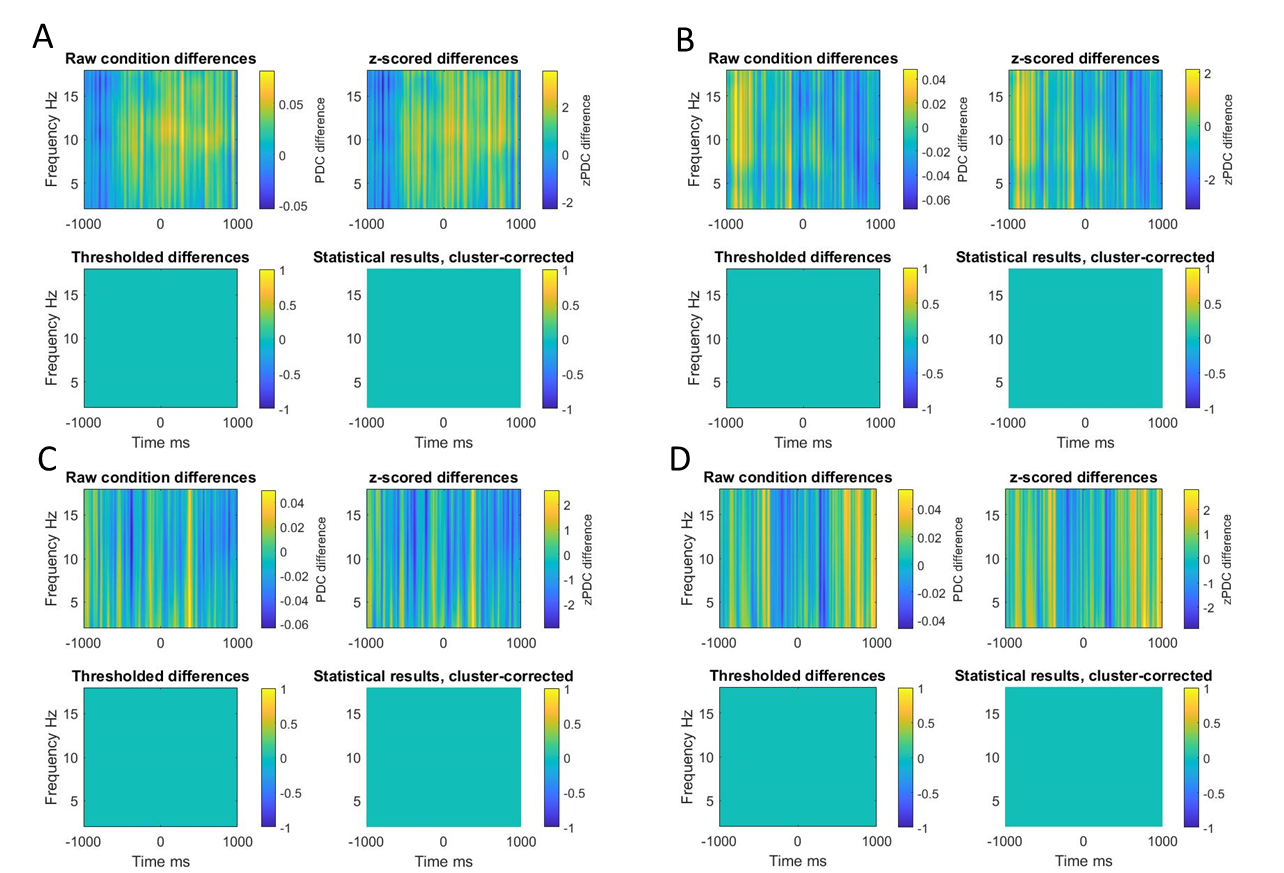

*Figure S12. Result of cluster-based permutation procedure for PDC over occipital electrodes. A) shows difference between infant sender/ adult receiver mutual vs non-mutual gaze onsets for I🡪A influences. B) shows difference between infant sender/ adult receiver mutual vs non-mutual for A🡪I influences. C) shows difference between adult sender/ infant receiver mutual vs non-mutual gaze onsets for I🡪A influences. D) shows difference between adult sender/ infant receiver mutual vs non-mutual gaze onsets for A🡪I influences (hotter, more yellow colours indicate more PLV for mutual vs non-mutual gaze). I🡪A means infant to adult influence. A🡪I means adult to infant influence.*

### 6. *Results of Bayes Factor analysis*

#### 6.1.  *Inter-brain non-event locked analysis – PLV and PDC*

|  | Theta | Alpha |
| --- | --- | --- |
| PLV | 0.19/ 0.49/ 5.31 | 0.16/ 0.74/ 6.35 |
| I🡪A PDC | 0.16/ 0.70/ 6.22 | 0.16/ 0.66/ 6.08 |
| A🡪I PDC | 0.18/ 0.50/ 5.37 | 0.19/ 0.48/ 5.25 |

*Table S1. Results of Bayes Factor analysis of inter-brain non-event locked PLV and PDC in theta and alpha. For each cell of the table the first number is the bf10 score, the second is the p value associated with the bf10 test and the third number is the bf01 score. These tests were conducted on average data over C3 and C4 electrodes in theta (3-6Hz) and alpha (6-9Hz).*

#### 6.2.  *Inter-brain event locked analysis – PLV and PDC*

|  | Theta | Alpha |
| --- | --- | --- |
| PLV | 0.24/ 0.34/ 4.24 | 0.37/ 0.17/ 2.69 |
| I🡪A PDC | 0.18/ 0.55/ 5.58 | 0.18/ 0.52/ 5.42 |
| A🡪I PDC | 0.18/ 0.58/ 5.71 | 0.18/ 0.57/ 5.67 |

*Table S2. Results of Bayes Factor analysis of inter-brain event locked PLV and PDC in theta and alpha. For each cell of the table the first number is the bf10 score, the second is the p value associated with the bf10 test and the third number is the bf01 score. These tests were conducted on average data over occipital electrodes in theta (3-6Hz) and alpha (6-9Hz).*

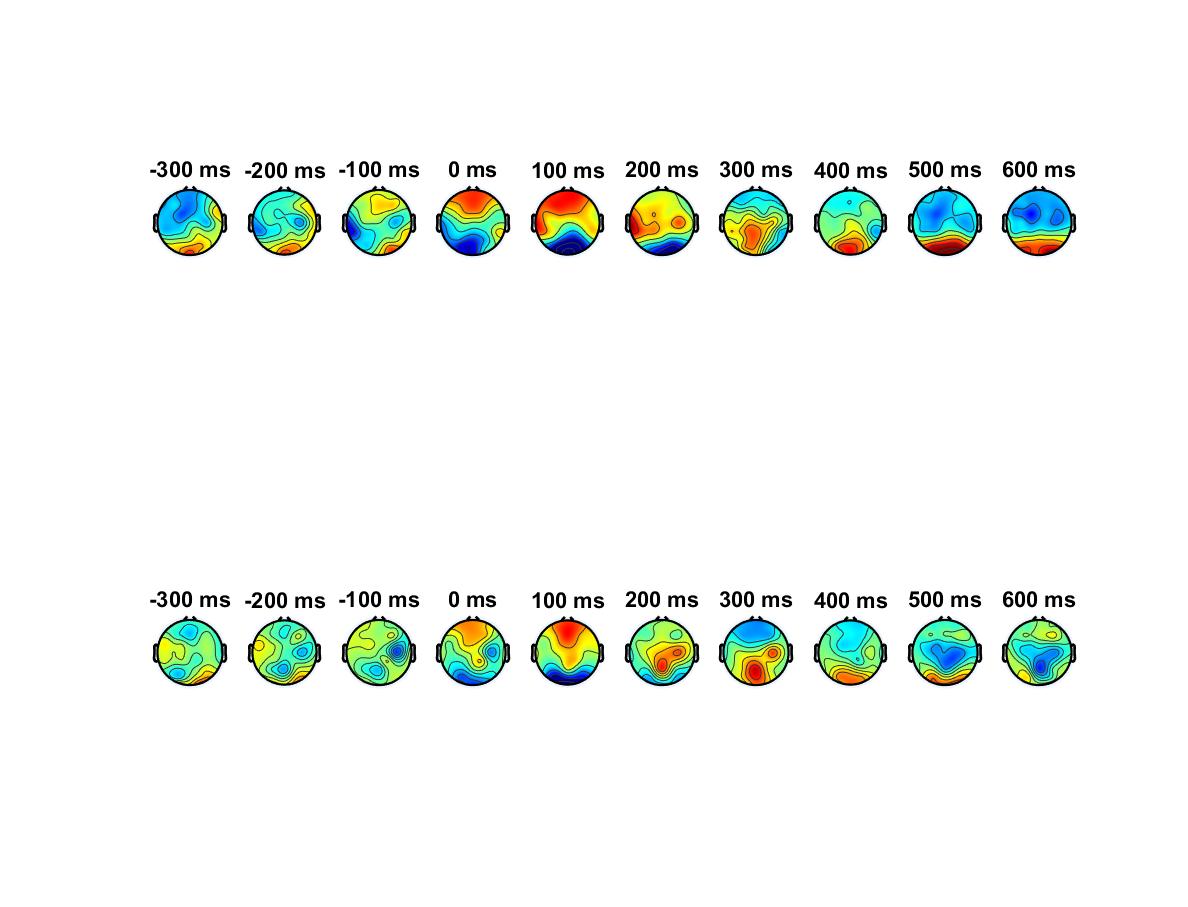

**A**

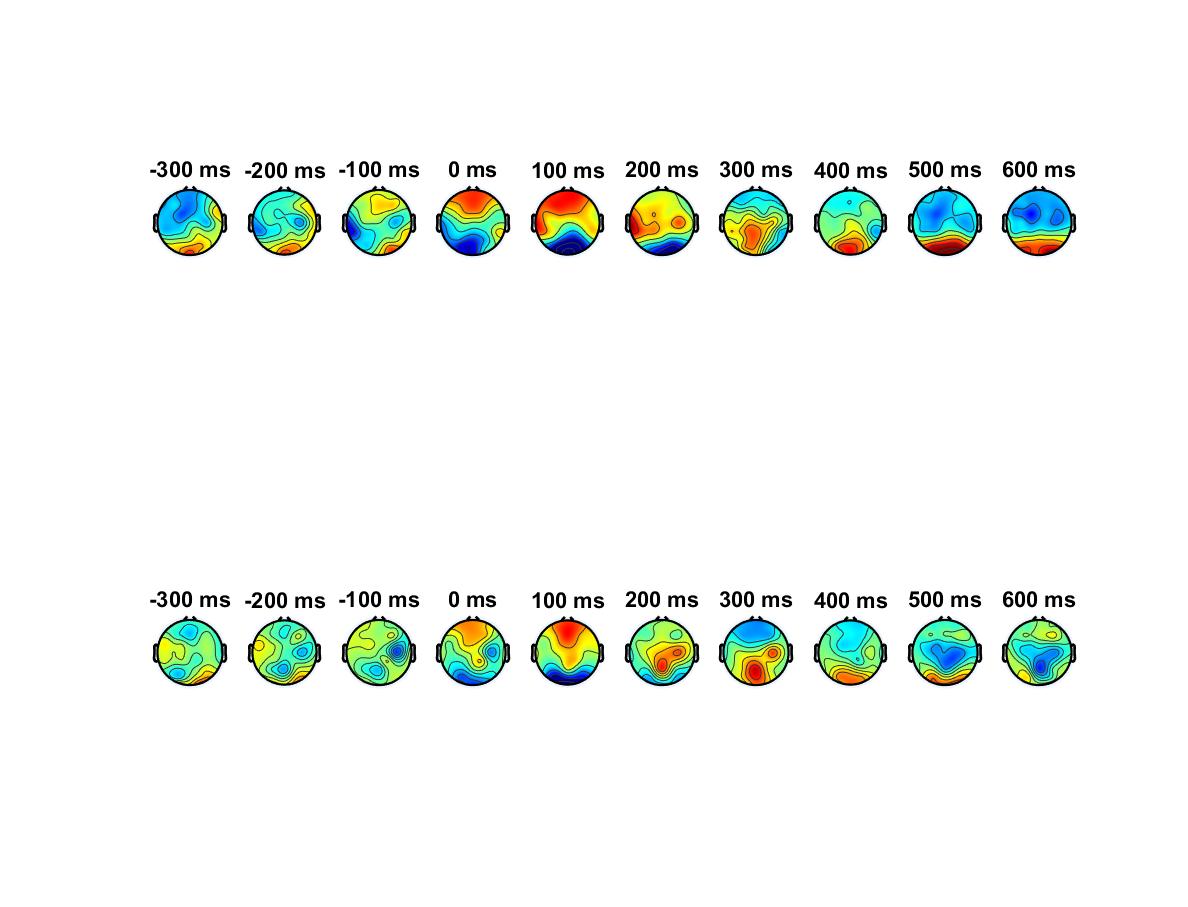

**B**

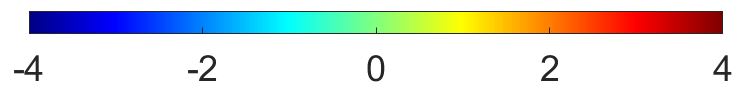

### 7. *Intra-brain analysis – ERP topoplots*

*Figure S13. Topographical distrbution of ERPs relative to infant sender mutual (A) and non-mutual (B) gaze onsets. Time 0 is onset of gaze.*

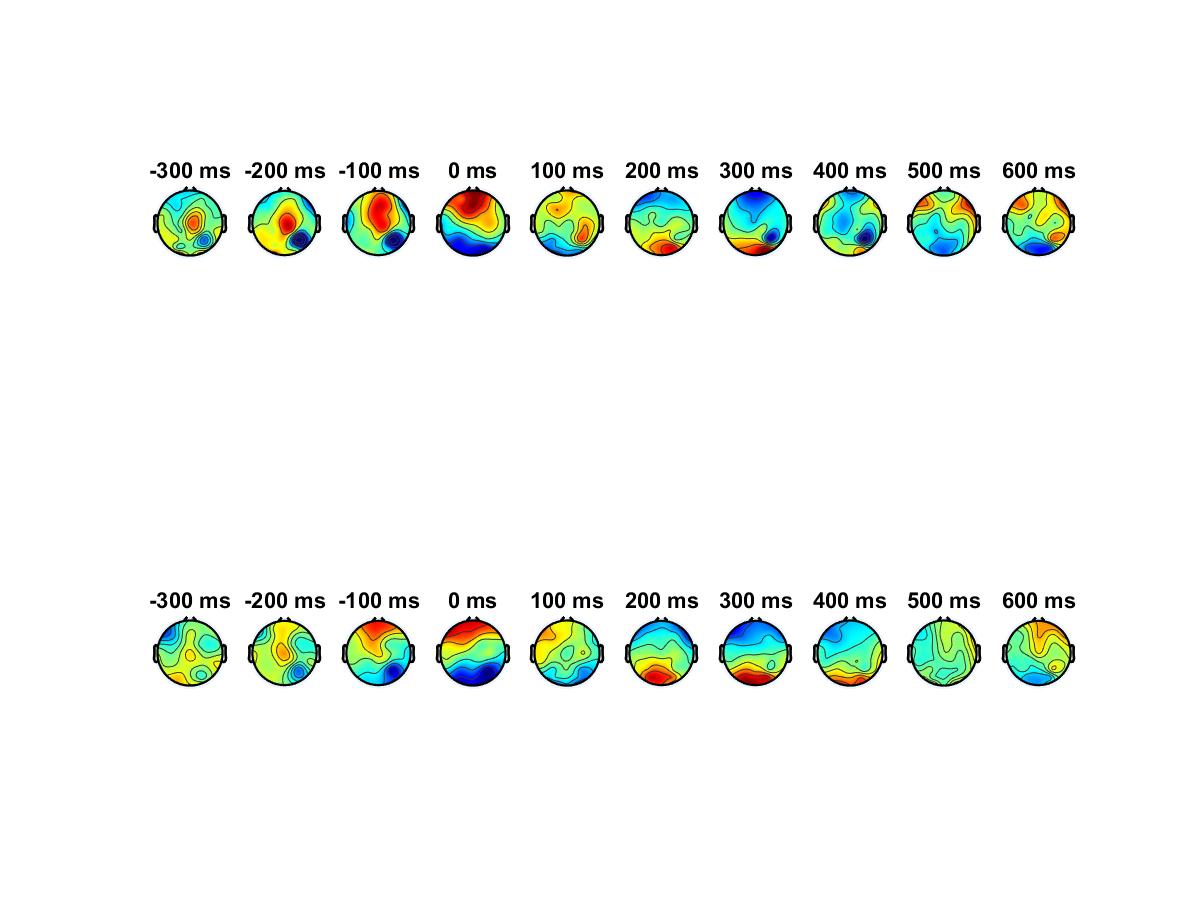

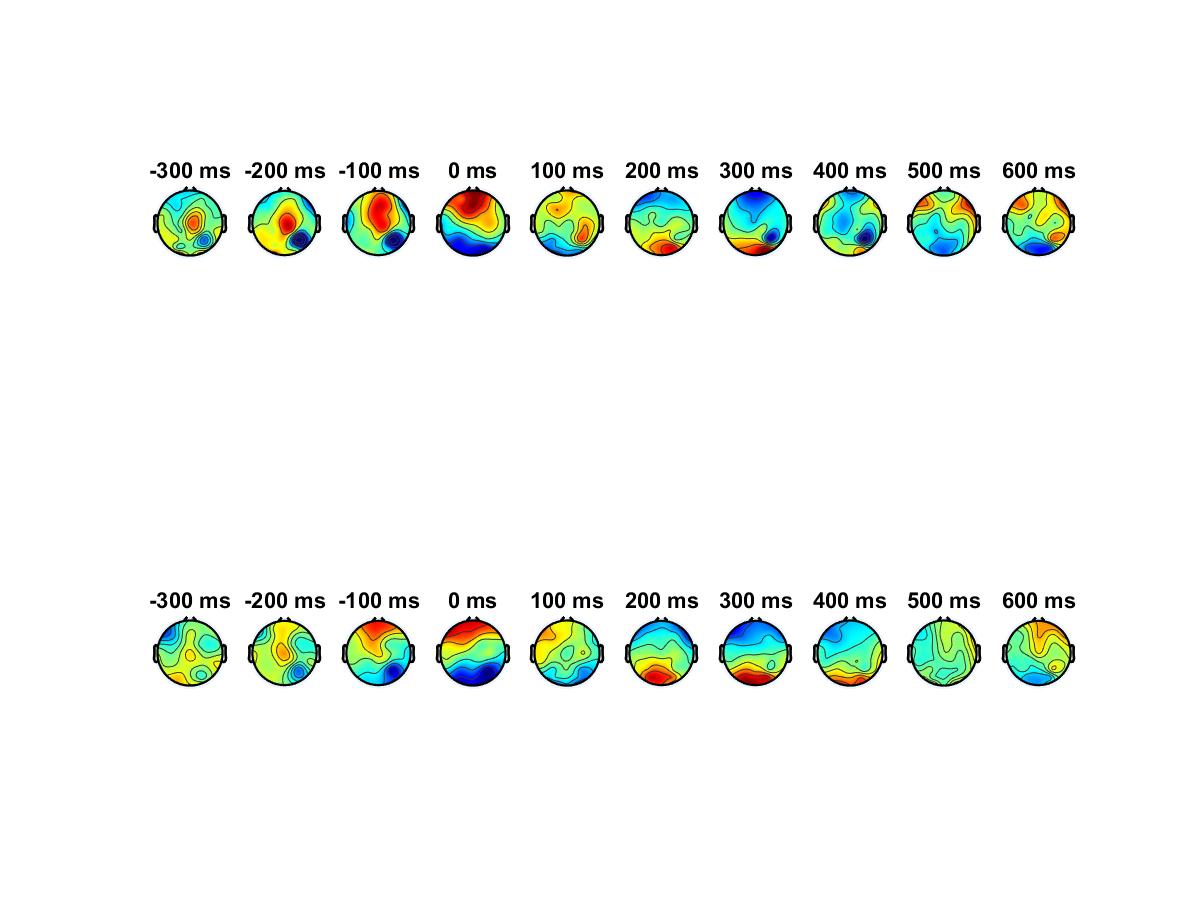

**B**

**A**

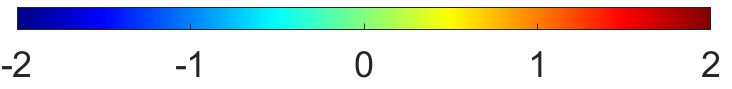

*Figure S14. Topographical distrbution of ERPs relative to infant sender mutual (A) and non-mutual (B) gaze onsets. Time 0 is onset of gaze.*

### 8. *Intra-brain analysis – ERP SEM*

In this section we report the standard error to the mean of single trail and group average (GA) ERP data for infant and adult, sender and recieiver, mutual and non-mutual gaze onsets.

|  | Sender | | | | | | Receiver | | | | | |
| --- | --- | --- | --- | --- | --- | --- | --- | --- | --- | --- | --- | --- |
|  | Mutual | | | Non-mutual | | | Mutual | | | Non-mutual | | |
| Sub no. | P1 | N290 | P400 | P1 | N290 | P400 | P1 | N290 | P400 | P1 | N290 | P400 |
| 1 | 1.98 | 2.04 | 1.79 | 2.59 | 6.15 | 3 | 2.15 | 1.84 | 2.11 | 1.31 | 1.23 | 1.19 |
| 2 | 1.81 | 3.39 | 1.54 | 2.26 | 2.17 | 1.78 | 2.15 | 2.21 | 2.07 | 1.94 | 2.48 | 1.72 |
| 3 | 5.05 | 10.82 | 11.54 | 5.64 | 5.82 | 4.56 | 4.93 | 8.43 | 4.99 | 4.05 | 5.04 | 8.1 |
| 4 | 2.2 | 2.81 | 1.44 | 2.21 | 3.44 | 2.68 | 1.18 | 1.25 | 1.04 | 1.66 | 2.03 | 1.75 |
| 5 | 4.4 | 1.82 | 6.82 | 7.85 | 7.43 | 3.72 | 2.5 | 3.59 | 3.16 | 3.77 | 3.24 | 2.81 |
| 6 | 1.46 | 2.99 | 2.44 | 1.09 | 1.62 | 0.96 | 1.65 | 1.07 | 1.46 | 1.1 | 1.22 | 0.99 |
| 7 | 4.95 | 4.79 | 3.14 | 2.26 | 6.98 | 6.33 | 5.09 | 3.09 | 3.53 | 9.05 | 3.73 | 5.92 |
| 8 | 1.55 | 1.66 | 1.58 | 2.03 | 2.44 | 0.65 | 4.6 | 2.24 | 2.6 | 1.31 | 1.33 | 1.05 |
| 9 | 2.57 | 2.38 | 2.54 | 3.76 | 3.12 | 2.82 | 4.58 | 4.9 | 6.2 | 2.45 | 2.75 | 2.63 |
| 10 | 1.82 | 1.74 | 1.65 | 2.62 | 3.93 | 3.73 | 3.74 | 2.13 | 2.99 | 2.6 | 1.59 | 1.96 |
| 11 | 7.63 | 5.73 | 3.72 | 6.62 | 7.9 | 7.56 | 8.42 | 3.19 | 2.5 | 3.68 | 3.31 | 1.95 |
| 12 | 2.35 | 2.94 | 2.76 | 4 | 3.97 | 3.82 | 5.26 | 4.46 | 2.65 | 2.37 | 1.62 | 2.17 |
| 13 | 1.01 | 1.14 | 0.95 | 0.56 | 1.53 | 4.88 | 1.56 | 1.42 | 1.75 | 0.83 | 0.93 | 0.74 |
| 14 | 1.11 | 1.07 | 1.28 | 3.78 | 4.89 | 7.63 | 5.32 | 4.83 | 2.35 | 1.19 | 1.5 | 1.47 |
| 15 | 2.87 | 3.24 | 1.8 | 3.22 | 3.36 | 2.75 | 2.9 | 2.82 | 1.78 | 2.14 | 3.62 | 1.82 |
| 16 | 8.83 | 6.79 | 8.61 | 3.04 | 5.1 | 4.27 | 2.24 | 5.12 | 2.92 | 3.33 | 5.45 | 4.37 |
| 17 | 2.5 | 2.31 | 2.1 | 1.45 | 3.15 | 1.92 | 3.3 | 3.26 | 2.51 | 1.48 | 1.35 | 1.21 |
| 18 | 2.29 | 2.59 | 2.16 | 3.01 | 2.47 | 3.67 | 2.2 | 1.97 | 2.2 | 2.16 | 2.06 | 1.73 |
| 19 | 2.3 | 2.77 | 2.36 | 2.34 | 3.51 | 1.7 | 2.39 | 5.35 | 3.53 | 3.58 | 4 | 2.91 |
| 20 | 3.11 | 3.83 | 3.4 | 2.14 | 3.45 | 2.28 | 1.59 | 1.9 | 1.67 | 2.04 | 2.11 | 1.66 |
| 21 | 2.71 | 2.09 | 1.89 | 3.87 | 4.14 | 3.58 | 3.05 | 2.84 | 2.55 | 2.19 | 1.7 | 1.83 |
| 22 | 1.83 | 1.6 | 1.45 | 2.41 | 4.4 | 3.45 | 1.4 | 1.37 | 1.37 | 1.11 | 0.82 | 1.04 |
| 23 | 14.35 | 16.39 | 19.69 | 6.31 | 10.15 | 6.45 | 9.49 | 16.96 | 7.15 | 15.05 | 11.42 | 15.51 |
| 24 | 3.6 | 2.94 | 2.44 | 7.09 | 3.22 | 1.98 | 5.55 | 4.46 | 4.06 | 2.93 | 2.58 | 3.25 |
| 25 | 2.5 | 2.25 | 2.56 | 3.8 | 3.2 | 3.13 | 3.27 | 3.78 | 3.62 | 1.65 | 1.61 | 1.52 |
| 26 | 2.44 | 2.32 | 1.98 | 3.21 | 4.3 | 4.52 | 2.72 | 3.66 | 3.18 | 2.4 | 2.74 | 2.54 |
| 27 | 3.67 | 3.79 | 2.83 | 6.46 | 4.33 | 4.24 | 1.91 | 2.98 | 3.25 | 2.35 | 2.68 | 1.94 |
| 28 | 3.28 | 3.24 | 3.16 | 4.13 | 4.17 | 2.86 | 4.14 | 4.96 | 2.65 | 4.57 | 3.01 | 2.76 |
| 29 | 6.52 | 18.05 | 13.79 | 37.63 | 39.29 | 19.09 | 21.88 | 53.31 | 3.76 | 8.6 | 8.95 | 12.09 |
| 30 | 2.37 | 3.05 | 2.44 | 4.28 | 4.37 | 2.89 | 5.78 | 10.94 | 7.02 | 2.45 | 3.07 | 2.02 |
| 31 | 2.23 | 7.21 | 4.89 | 6.51 | 6.19 | 3.3 | 3.5 | 4 | 4.62 | 1.81 | 1.97 | 2.02 |
| 32 | 11.97 | 12.97 | 5.53 | 5.85 | 8.49 | 4.76 | 4.28 | 5.87 | 4.46 | 7.04 | 7.38 | 3.38 |
| 33 | 1.84 | 2.05 | 1.97 | 5.54 | 5.53 | 3.14 | 3.35 | 2.93 | 3.3 | 1.94 | 2.43 | 2.4 |
| 34 | 1.92 | 2.36 | 1.97 | 2.81 | 3.38 | 1.88 | 2.74 | 1.47 | 1.86 | 1.15 | 1.04 | 0.97 |
| 35 | 4.86 | 6.43 | 4.08 | 3.31 | 3.73 | 4.56 | 4.33 | 4.98 | 4.38 | 5.29 | 5.87 | 4.24 |
| 36 | 3.1 | 3.05 | 2 | 3 | 3.77 | 2.88 | 3.54 | 4.41 | 3.28 | 3.8 | 3.03 | 2.62 |
| 37 | 2.35 | 3.2 | 4.43 | 4.71 | 1.75 | 2.18 | 3.87 | 8.69 | 3.87 | 3.42 | 2.19 | 2.03 |
| 38 | 9.08 | 2.2 | 2.43 | 5.92 | 3 | 3.95 | 6.16 | 3.76 | 3.77 | 1.85 | 2.5 | 2.19 |
| 39 | 1.99 | 2.42 | 1.82 | 2.17 | 4.42 | 6.13 | 2.62 | 2.23 | 4.29 | 2.19 | 2.22 | 2.05 |
| 40 | 2.62 | 4.82 | 2.25 | 2.69 | 10.19 | 4.85 | 2.45 | 3.75 | 2.98 | 5 | 4.61 | 3.73 |
| 41 | 8.1 | 9.09 | 5.57 | 14.4 | 9.59 | 13.46 | 11.24 | 15.98 | 13.22 | 2.28 | 2.4 | 2.67 |
| 42 | 3.06 | 4.02 | 3.87 | 3.81 | 3.68 | 4.82 | 2.99 | 2.96 | 3.34 | 1.8 | 1.4 | 1.32 |
| 43 | 2.3 | 2.17 | 2.51 | 2.31 | 1.87 | 2.66 | 2.18 | 1.51 | 1.8 | 1.18 | 1.25 | 1.06 |
| 44 | 3 | 2.42 | 2.98 | 5.13 | 3.67 | 1.95 | 2.63 | 3.79 | 2.61 | 1.41 | 1.39 | 1.61 |
| 45 | 10.76 | 10.8 | 13.12 | 4.05 | 5.37 | 5.21 | 7.18 | 5.6 | 7.59 | 4.71 | 4.46 | 4.41 |
| 46 | 10.23 | 20.42 | 24.94 | 120.06 | 99.47 | 21.62 | 42.25 | 47.59 | 44.93 | 24.03 | 19.19 | 16.44 |
| 47 | 2.26 | 4.19 | 2.47 | 3.46 | 7.14 | 4.18 | 2.96 | 2.02 | 2.34 | 1.28 | 1.95 | 1.13 |
| 48 | 3.86 | 3.43 | 2.58 | 3.81 | 3.85 | 1.52 | 3.09 | 4.3 | 1.64 | 4.42 | 8.59 | 4.53 |
| 49 | 13.84 | 15.65 | 7.94 | 4.33 | 6.03 | 5.47 | 5.26 | 4.96 | 5.87 | 13.61 | 21.64 | 12.72 |
| 50 | 1.39 | 2.28 | 1.67 | 2.58 | 1.82 | 1.98 | 2.48 | 2.52 | 2 | 2.64 | 4.15 | 2.35 |
| 51 | 2.22 | 3.39 | 1.99 | 4.15 | 3.1 | 3.2 | 2.56 | 1.61 | 1.85 | 2.45 | 1.85 | 1.8 |
| 52 | 36.87 | 34.8 | 37.67 | 8.41 | 6.63 | 7.99 | 15.59 | 17.34 | 19.96 | 10.8 | 21.13 | 8.14 |
| 53 | 12.28 | 8.99 | 13.67 | 10.76 | 2.85 | 13.6 | 14.03 | 17.64 | 30.24 | 8 | 5.63 | 5.71 |
| GA | 1.26 | 2.23 | 3.32 | 5.22 | 2.12 | 2.41 | 1.26 | 0.9 | 3.32 | 0.99 | 2.12 | 1 |

*Table S3. Results of standard error of the mean tests of infant ERP data. Note that removing outliers did not impact the significance of any of the tests, i.e. no tests that were not significant before became significant after removing outlier data a re-testing.*

|  | Sender | | | | | | Receiver | | | | | |
| --- | --- | --- | --- | --- | --- | --- | --- | --- | --- | --- | --- | --- |
|  | Mutual | | | Non-mutual | | | Mutual | | | Non-mutual | | |
| Sub no. | P1 | N170 | P300 | P1 | N170 | P300 | P1 | N170 | P300 | P1 | N170 | P300 |
| 1 | 284.78 | 291.87 | 196.88 | 359.57 | 227.26 | 338.18 | 307.82 | 198.98 | 269.4 | 273.98 | 108.67 | 175.12 |
| 2 | 111.08 | 121.85 | 109.46 | 133.38 | 105.24 | 196.23 | 106.21 | 162.83 | 115.62 | 206.06 | 177.56 | 164.46 |
| 3 | 1.25 | 1.29 | 1.24 | 2.14 | 1.78 | 2.28 | 1.86 | 1.62 | 1.37 | 1.22 | 1.53 | 1.12 |
| 4 | 0.62 | 0.7 | 0.69 | 0.57 | 0.9 | 1.22 | 0.41 | 0.45 | 0.49 | 0.49 | 0.64 | 0.61 |
| 5 | 1.83 | 2.81 | 1.89 | 3.22 | 4.14 | 2.71 | 3.42 | 2.96 | 2.53 | 1.93 | 2.93 | 2.51 |
| 6 | 0.48 | 0.64 | 0.42 | 0.49 | 0.49 | 0.42 | 0.7 | 0.76 | 0.41 | 0.56 | 0.74 | 0.46 |
| 7 | 1.22 | 0.87 | 1.05 | 1.71 | 2.48 | 1.39 | 1.94 | 2.46 | 1.59 | 1.83 | 2.19 | 2.16 |
| 8 | 4.27 | 3.05 | 3.83 | 6.89 | 10.03 | 9.64 | 13.5 | 6.2 | 4.52 | 1.74 | 1.86 | 2.29 |
| 9 | 1.1 | 1.23 | 1.1 | 1.35 | 1.21 | 1.15 | 0.91 | 1.08 | 0.99 | 1.68 | 1.59 | 1.25 |
| 10 | 2.34 | 1.92 | 2.97 | 4.45 | 4.05 | 4.82 | 3.85 | 4.67 | 3.27 | 2.27 | 2.28 | 2.74 |
| 11 | 4.52 | 1.72 | 3.52 | 3.29 | 3.82 | 3.28 | 6.72 | 6.03 | 5.02 | 5.02 | 1.23 | 2.88 |
| 12 | 1.01 | 0.96 | 0.94 | 2.12 | 3.59 | 1.74 | 1.21 | 1.67 | 2.55 | 1.15 | 1.06 | 0.96 |
| 13 | 1.94 | 1.09 | 1.6 | 4.08 | 3.94 | 5.91 | 1.69 | 1.28 | 1.91 | 0.77 | 0.76 | 0.8 |
| 14 | 0.93 | 1.23 | 0.82 | 4.05 | 3.9 | 1.15 | 2.67 | 3.42 | 2.54 | 0.89 | 1.04 | 0.93 |
| 15 | 1.1 | 1.24 | 0.94 | 2.95 | 2.62 | 1.19 | 1.17 | 1.19 | 1.44 | 0.96 | 1.41 | 0.95 |
| 16 | 1.23 | 1.07 | 0.9 | 1.86 | 1.34 | 1.22 | 2.6 | 2.25 | 1.71 | 1.12 | 1.46 | 1.33 |
| 17 | 1.16 | 1.25 | 1.24 | 1.06 | 1.09 | 1.28 | 1.61 | 1.81 | 1.38 | 1.48 | 1.38 | 0.97 |
| 18 | 5.92 | 3.34 | 5.41 | 1.36 | 1.65 | 2.11 | 8.54 | 4.28 | 4.35 | 0.77 | 1.1 | 0.77 |
| 19 | 0.66 | 0.5 | 0.62 | 6.28 | 7.91 | 23.3 | 2.59 | 1.84 | 1.43 | 7.73 | 5.22 | 4.38 |
| 20 | 1.45 | 2.04 | 1.44 | 0.79 | 0.67 | 1.44 | 1.24 | 1.27 | 0.92 | 0.72 | 1.21 | 0.8 |
| 21 | 2.35 | 3.29 | 2.29 | 1.53 | 1.72 | 2.43 | 1.69 | 1.52 | 1.82 | 1.14 | 1.13 | 0.92 |
| 22 | 2.5 | 2.86 | 2.09 | 1.32 | 1.31 | 1.54 | 1.75 | 3.24 | 1.68 | 1.11 | 0.71 | 0.81 |
| 23 | 1.32 | 2.34 | 1.56 | 4.52 | 4.46 | 3.75 | 3.95 | 6.49 | 1.78 | 2.58 | 2.05 | 2.03 |
| 24 | 3.66 | 2.87 | 2.58 | 2.45 | 5.31 | 5.23 | 5.72 | 3.79 | 3.72 | 2.24 | 4.37 | 2.93 |
| 25 | 1.27 | 1.26 | 1.3 | 0.62 | 0.94 | 0.23 | 1.84 | 2.05 | 1.81 | 6.37 | 3.23 | 2.34 |
| 26 | 4.38 | 2.95 | 2.5 | 2.62 | 1.54 | 2.14 | 2.37 | 2.57 | 2.02 | 0.73 | 0.82 | 0.73 |
| 27 | 1.28 | 1.35 | 1.52 | 2.92 | 3.95 | 5.87 | 1.78 | 1.63 | 1.97 | 3.29 | 2.31 | 2.85 |
| 28 | 1.21 | 1.13 | 1.04 | 2.43 | 2.54 | 2.95 | 2.18 | 2.14 | 3.01 | 0.89 | 1.03 | 0.77 |
| 29 | 2.88 | 2.67 | 1.94 | 1.1 | 1.86 | 1.26 | 5.16 | 0.35 | 4.02 | 1.54 | 1.2 | 1.68 |
| 30 | 1.56 | 3.74 | 4.32 | 9.02 | 12.08 | 6.15 | 4.03 | 2.08 | 2.79 | 4.53 | 2.02 | 2.72 |
| 31 | 4.57 | 1.82 | 4.4 | 1.13 | 1.15 | 0.96 | 2.75 | 1.58 | 3.69 | 0.97 | 1.06 | 0.94 |
| 32 | 3.16 | 3.27 | 2.6 | 2.28 | 0.96 | 2.14 | 5.56 | 3.65 | 3.98 | 1.65 | 1.99 | 1.87 |
| 33 | 0.9 | 1.21 | 0.86 | 2.82 | 2.77 | 2.72 | 1.67 | 1.31 | 1.58 | 2.05 | 1.35 | 1.37 |
| 34 | 1.1 | 1.14 | 0.91 | 3.42 | 4.03 | 1.94 | 1.15 | 1.05 | 1.04 | 1.23 | 1.45 | 0.91 |
| 35 | 1.49 | 1.31 | 1.57 | 1.44 | 1.4 | 1.53 | 1.48 | 1.91 | 2.04 | 0.62 | 0.61 | 0.71 |
| 36 | 0.46 | 0.62 | 0.68 | 2.16 | 1.94 | 2.3 | 0.78 | 1.55 | 0.49 | 1.16 | 1.07 | 0.75 |
| 37 | 1.3 | 1.52 | 1.01 | 0.6 | 0.38 | 0.46 | 1.46 | 1.15 | 1.19 | 0.4 | 0.37 | 0.39 |
| 38 | 0.9 | 0.94 | 1.02 | 0.85 | 0.93 | 0.93 | 1.64 | 1.8 | 1.05 | 1.04 | 1.42 | 1.59 |
| 39 | 1.69 | 1.15 | 1.5 | 2.84 | 1.56 | 1.41 | 1.58 | 2.28 | 1.56 | 1.36 | 1.24 | 1.09 |
| 40 | 4.23 | 2.62 | 3.22 | 1.75 | 1.95 | 1.46 | 5.92 | 4.09 | 4.2 | 0.81 | 0.8 | 0.83 |
| 41 | 1.57 | 1.59 | 1.37 | 29.09 | 13.31 | 10.47 | 4.59 | 2.41 | 1.77 | 9.36 | 5.36 | 9.08 |
| 42 | 7.08 | 10.04 | 8.6 | 4.14 | 4.16 | 1.94 | 11.06 | 12.77 | 11.81 | 1.35 | 1.2 | 0.94 |
| 43 | 1.09 | 1.42 | 1.07 | 28.07 | 17.53 | 29.14 | 1.3 | 1.38 | 1.35 | 6.61 | 6.48 | 6.07 |
| 44 | 1.72 | 1.2 | 1.16 | 1.57 | 0.92 | 1.94 | 1.48 | 1 | 0.83 | 1.14 | 0.94 | 0.98 |
| 45 | 1.66 | 3.78 | 2 | 1.73 | 1.61 | 1.43 | 7.08 | 5.67 | 6.15 | 0.84 | 0.77 | 0.54 |
| 46 | 3.18 | 4.36 | 6.74 | 8.67 | 5.45 | 2.12 | 4.98 | 3.02 | 3.41 | 1.93 | 1.82 | 1.36 |
| 47 | 41.98 | 56.28 | 46.06 | 2.91 | 3.32 | 0.76 | 64.97 | 73.52 | 90.82 | 2.05 | 1.46 | 2.75 |
| 48 | 8.23 | 2.69 | 8.32 | 95.69 | 48.11 | 72.56 | 3.73 | 3.45 | 2.96 | 35.5 | 44.77 | 55.47 |
| 49 | 2.55 | 2.6 | 1.86 | 9.46 | 5.74 | 7.92 | 1.77 | 2.02 | 2.3 | 2.69 | 2.33 | 3.06 |
| 50 | 1.9 | 1.13 | 1.61 | 3.12 | 3.58 | 4.35 | 2.42 | 1.31 | 1 | 2.07 | 10.36 | 1.66 |
| 51 | 209.04 | 131.15 | 200.34 | 6.81 | 2.33 | 5.28 | 569.93 | 958.7 | 624.33 | 2.79 | 5.34 | 4.01 |
| 52 | 57.3 | 23.1 | 42.75 | 1887.85 | 486.15 | 2090.58 | 24.53 | 10.17 | 15.13 | 446.77 | 1148.22 | 1181.83 |
| 53 | 1.52 | 0.74 | 1.41 | 16.55 | 7.24 | 17.88 | 0.35 | 0.92 | 4.59 | 3.74 | 5.67 | 1.3 |
| GA | 11.13 | 68.16 | 7.7 | 28.94 | 6.17 | 41.34 | 15.79 | 17.11 | 11.63 | 19.6 | 15.8 | 37.39 |

*Table S4. Results of standard error of the mean tests of adult ERP data. Note that removing outliers did not impact the significance of any of the tests, i.e. no tests that were not significant before became significant after removing outlier data a retesting*

#### **References**

Chaumon, M., Bishop, D. V., & Busch, N. A. (2015). A practical guide to the selection of independent components of the electroencephalogram for artifact correction. *Journal of neuroscience methods*, *250*, 47-63.

Muthukumaraswamy, S. D., & Singh, K. D. (2011). A cautionary note on the interpretation of phase-locking estimates with concurrent changes in power. *Clinical neurophysiology: official journal of the International Federation of Clinical Neurophysiology*, *122*(11), 2324-2325.

Haresign, I. M., Phillips, E., Whitehorn, M., Noreika, V., Jones, E. J. H., Leong, V., & Wass, S. V. (2021). Automatic classification of ICA components from infant EEG using MARA. *Developmental cognitive neuroscience*, *52*, 101024.
